# Failures of nerve regeneration caused by aging or chronic denervation are rescued by restoring Schwann cell c-Jun

**DOI:** 10.1101/2020.10.06.327957

**Authors:** Laura J. Wagstaff, Jose A. Gomez-Sanchez, Shaline V. Fazal, Georg W. Otto, Alastair M. Kilpatrick, Kirolos Michael, Liam Y.N. Wong, Ki H. Ma, Mark Turmaine, John Svaren, Tessa Gordon, Peter Arthur-Farraj, Sergio Velasco-Aviles, Hugo Cabedo, Cristina Benito, Rhona Mirsky, Kristjan R Jessen

## Abstract

After nerve injury, myelin and Remak Schwann cells reprogram to repair cells specialized for regeneration. Normally providing strong regenerative support, these cells fail in aging animals, and during the chronic denervation that results from the slow growth of axons. This impairs axonal regeneration and causes a significant clinical problem. In mice, we find that repair cells express reduced c-Jun protein as the regenerative support provided by these cells declines in aging animals and during chronic denervation. In both cases, genetically restoring Schwann cell c-Jun levels restores regeneration to that in controls. We identify potential gene candidates mediating this effect and implicate Shh in the control of Schwann cell c-Jun levels. This establishes that a common mechanism, reduced c-Jun in Schwann cells, regulates the success and failure of nerve repair both during aging and chronic denervation. This provides a molecular framework for addressing important clinical problems, and suggests molecular pathways that can be targeted to promote repair in the PNS.

## INTRODUCTION

Among mammalian systems, peripheral nerve is often hailed as a prime example of a tissue with a striking regenerative potential. Nerve injury triggers the reprograming of myelin and non-myelin (Remak) Schwann cells to adopt a repair Schwann cell phenotype specialized to support regeneration, and injured neurons activate a gene program that facilitates axon growth. Yet, paradoxically, the clinical outcome of nerve injuries remains poor, and nerve damage constitutes a significant clinical and economic burden. Remarkably, treatment of nerve injuries has not advanced significantly for decades (Furey et al. 2007; Jonsson et al. 2013; reviewed in Fu and Gordon 1995; Boyd and Gordon 2003a; Hoke 2006; Allodi et al. 2012; Scheib and Hoke 2013; Doron-Mandel et al. 2015; Jessen and Mirsky 2016; Fawcett and Verhaagen 2018; Jessen and Arthur-Farraj 2019).

The question of why a potentially regenerative tissue fails to respond effectively to injury and ensure clinical recovery is important both for promoting nerve repair, and also more generally. A number of other systems with experimentally established regenerative capacity, e.g. skin, heart and pancreatic islets, also fail to show clinically useful regenerative response to tissue damage (Cohen and Melton, 2011; Eguizabal et al. 2013; Jessen et al. 2015).

In the case of peripheral nerves, recent work has highlighted two important factors that prevent full expression of their regenerative potential. One is the age of the animal at the time of injury, increasing age resulting in a marked decrease in regeneration. The other is the adverse effect of chronic denervation on the nerve distal to injury, since this tissue gradually loses the capacity to support axon growth as it lies denervated during the often extensive time it takes regenerating axons to reach their targets. These two problems turn out to involve a common factor, namely a repair Schwann cell failure, since both during aging and chronic denervation, the denervated Schwann cells in the distal stump undergo molecular and morphological changes that result in a striking functional deterioration of these important drivers of axonal regeneration (reviewed in Verdu et al 2000; Sulaiman and Gordon 2009; Painter 2017; Jessen and Mirsky 2019).

In the present work, we have tested whether the dysfunction of repair Schwann cells in these two apparently unrelated situations relates to a common factor, namely a failure to activate or maintain high levels of the transcription factor c-Jun. That this might be so, is based on our previous finding that c-Jun, which is upregulated in Schwann cells in injured nerves, is a global amplifier of the repair Schwann cell phenotype (Arthur-Farraj et al. 2012; reviewed in Jessen and Mirsky 2016; 2019; Jessen and Arthur-Farraj 2019), and on subsequent findings showing that enhanced Schwann cell c-Jun promotes regeneration, both through nerve grafts and *in vitro* (Arthur-Farraj et al. 2012; Huang et al. 2015; 2019).

The age-dependent decline in regenerative capacity of human and animal nerves is well established (Pestronk et al. 1980; Tanaka and Webster 1991; Tanaka et al. 1992; Graciarena et al. 2014; reviewed in Vaughan 1992; Verdú et al. 2000; Ruijs et al. 2005). This is associated with a reduced initial inflammatory response followed by enhanced chronic inflammation (Scheib and Höke 2016; Büttner et al. 2018). Interestingly, diminished regeneration is not caused by age-dependent changes in neurons. Rather, aging results in subdued activation of the repair Schwann cell phenotype after injury, including reduced c-Jun expression, resulting in regeneration failure (Painter et al. 2014; reviewed in Painter 2017).

The other major barrier to repair that we consider here is caused by long-term denervation of nerves distal to injury. This is an important issue in human nerve regeneration (Ruiis et al. 2005), and has been studied in some detail in rats, revealing that chronic denervation results in reduced expression of repair-associated genes including GDNF, BDNF, NT-3 and p75NTR, accompanied by a dramatic reduction in the ability of denervated distal stumps to support regeneration even of freshly transected axons (Fu and Gordon 1995; You et al. 1997; Sulaiman and Gordon 2000; Höke et al. 2002; Michalski et al. 2008; Eggers et al. 2010). There is direct evidence for a comparable deterioration of repair cells and repair capacity in during chronic denervation of human nerves (Wilcox et al. 2020; reviewed in Ruijs et al. 2005). Chronic denervation also results in reduced repair cell numbers and shortening of repair cells (Benito et al. 2017; Gomez-Sanchez et al. 2017; reviewed in Jessen and Mirsky 2019). Thus, the repair phenotype is not stable, but fades with time after injury, thereby contributing to the poor outcome after nerve damage in humans.

Schwann cell reprograming after nerve injury involves up-regulation of trophic factors and cytokines, activation of EMT genes, and myelin autophagy for myelin clearance and down-regulation of myelin genes (Brushart et al. 2013; Arthur-Farraj et al. 2017; Clements et al. 2017; reviewed in Gröthe et al. 2006; Chen et al 2007; Gambarotta et al. 2013; Glenn and Talbot 2013; Jessen and Mirsky 2016; Boerboom et al. 2017; Jessen and Arthur-Farraj 2019; Nocera and Jacob 2020). Myelin and Remak Schwann cells also increase in length by two-to-three fold and often branch as they convert to repair cells and form regeneration tracks, Bungner bands, that guide regenerating axons (Gomez-Sanchez et al. 2017). The molecular signals involved in the decline of these repair-supportive features during aging and chronic denervation have not been known.

The transcription factor c-Jun regulates the reprograming of myelin and Remak cells to repair cells by accelerating the extinction of myelin genes, promoting myelin breakdown, and by amplifying the up-regulation of a broad spectrum of repair-supportive features, including the expression of trophic factors. Accordingly, genetic removal of c-Jun from Schwann cells results in functionally impaired repair cells and regeneration failure (Arthur-Farraj et al. 2012; Fontana et al., 2012; reviewed in Jessen and Arthur-Farraj 2019).

Here we provide evidence that a common molecular mechanism, the dysregulation of c-Jun in Schwann cells, is central to two major categories of regeneration failure in the PNS. The high levels of Schwann cell c-Jun triggered by nerve injury in young animals are not achieved in older ones, and, irrespective of age, the elevated c-Jun expression seen after injury steadily decreases during long-term denervation. Importantly we show that in both models of regeneration failure, genetically restoring Schwann cell c-Jun levels in vivo also restores regeneration rates to that in controls. By establishing c-Jun as an important regulator of the success and failure of nerve repair during aging and chronic denervation this observation provides a common molecular framework for addressing an important clinical problem, and suggests molecular pathways that can be targeted to promote repair in the PNS.

## RESULTS

### IN AGING ANIMALS, MAINTAINING C-JUN LEVELS IN SCHWANN CELLS REVERSES AGE-RELATED DEFECTS IN NERVE REGENERATION

Age-dependent failure of nerve regeneration is accompanied by subdued elevation of c-Jun, a major regulator of the repair cell phenotype (Painter et al. 2014). To test whether this reduction in c-Jun controls the reduced capacity of these cells to support axon growth, we first compared c-Jun upregulation in young and older WT mice (Fig.1 A, B). Four days after transection, c-Jun protein levels in the distal nerve stump in aged mice (8-10 month) were found to be ~ 50% lower than in young (6-8 week) mice.

**Figure 1:**
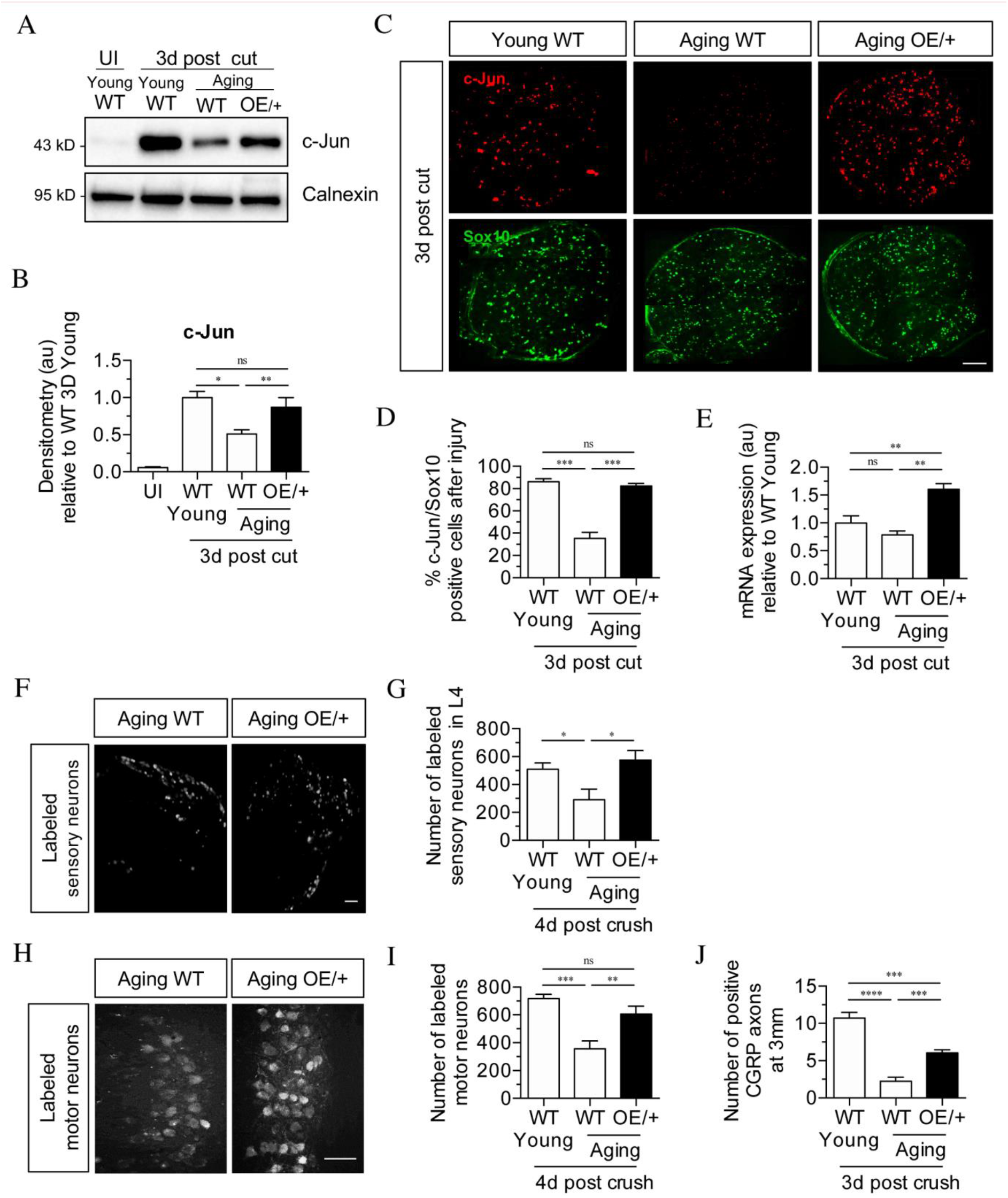
Restoring Schwann cell c-Jun protein reverses the age-related decline in nerve regeneration. (A) Representative Western blots of c-Jun in young and aging WT nerves and aging c-Jun OE/+ nerves 3 days post injury. Quantification by densitometry is in (B). c-Jun up-regulation is impaired in WT aged nerves but restored in aged c-Jun OE/+ nerves. Data normalised to young WT 3 days post cut, *p<0.05, **p<0.005, ns, non-significant. Young UI WT n=6, n=5 for all other experimental groups. (C) Representative images showing immunofluorescence of c-Jun in double labeling with Schwann cell nuclear marker Sox10 in sections of the distal nerve stump in young and aging WT and aging c-Jun OE/+ mice 3 days post cut. Quantification by cell counting is in (D). In aging WT Schwann cells, c-Jun is reduced, but elevated to youthful levels in aging c-Jun OE/+ Schwann cells. ***p<0.001, ns, nonsignificant. n=3 for each experimental group. (E) RTqPCR analysis of 3 day cut nerves. Data normalised to young WT 3 days post cut. **p<0.005, ns, non-significant, n=4 (F) Representative images showing Fluorogold-labeled sensory neurons in L4 DRGs of aging WT and c-Jun OE/+ mice 1 week post back-filling following a 3 day crush injury. Quantification by cell counting is in (G). There is an age-related decrease in back-filled neurons in WT samples (p=0.0309), while the high number of regenerating neurons in young WT mice is maintained in aging c-Jun OE/+ DRGs (p=0.0211). Unpaired Student’s t-test. Young WT n=6, Aging WT n=5, Aging c-Jun OE/+ n=6. (H) Representative images of Fluorogold-labeled motor neurons in aging WT and c-Jun OE/+ mice 1 week post back-filling following a 3 day crush injury. Quantification is in (I). Counts of labeled motor neurons mirror those of sensory neurons since in WT mice, but not in c-Jun OE/+ mice, the number of back-labeled motor neurons decreases with age. ***p<0.001, **p<0.005, n=6 for all experimental groups. (J) Counts of calcitonin gene-related peptide (CGRP)+ regenerating axons 3mm from crush injury of the sciatic nerve of young and aged WT mice, and of aging c-Jun OE/+ mice. ****p<0.0001, ***p<0.001. Young WT n=5, aging WT and c-Jun OE/+ n=6. All numerical data are analysed by one-way ANOVA with Tukey’s multiple comparison test and represented as means ± SEM. All scale bars: 100μm.

To determine the functional significance of this, we studied P0Cre +/-; c-Jun f/+ mice (referred to as c-Jun OE/+ mice), which we generated previously (Fazal et al. 2017). In these mice, c-Jun levels are enhanced in Schwann cells only. In Western blots of uninjured adult sciatic nerves of c-Jun OE/+ mice, c-Jun is elevated about seven fold compared to WT. While there is a modest reduction in myelin thickness, nerve architecture and Schwann cell morphology are normal (Fazal et al. 2017). We found that in c-Jun OE/+ mice, the age-dependent decline in c-Jun protein levels after sciatic nerve cut was prevented, and that c-Jun levels in the distal stump of young WT and aging c-Jun OE/+ mice were similar by Western blots 4 days after cut (Fig.1 A, B). This was confirmed in immunofluorescence experiments on 3 day cut nerves, using Sox10 antibodies to selectively identify Schwann cell nuclei, and c-Jun antibodies (Fig.1C, D). In WT mice, older nerves contained fewer c-Jun positive Schwann cell nuclei and the labelling of the c-Jun positive nuclei was weaker, compared to young nerves. In aged c-Jun OE/+ nerves nuclear c-Jun was restored to levels similar to those in young WT nerves. At the mRNA level, a non-significant trend towards lower c-Jun expression was seen in 3 day cut nerves of aged WT nerves, while there was a significant elevation of c-Jun mRNA in cut c-Jun OE/+ nerves as expected (Fig.1 E).

Regeneration in young and aged WT mice and aged c-Jun OE/+ mice was compared using neurone back-filling, a method that provides an optimal measure of regenerative capacity by determining the number of neuronal cell bodies have regenerated axons through a nerve at a measured distance distal to injury (Novikova et al.1997; Boyd and Gordon 2003b; Catapano et al. 2016). Four days after sciatic nerve crush, retrograde tracer was applied to the distal stump 7 mm from the crush site. Seven days later, the animals were sacrificed and the number of back-filled DRG and spinal cord motor neurons were counted. The results were comparable for both neuronal populations (Fig.1 F-J). The number of neurons regenerating through the distal stump of aged WT mice was reduced by about 50% compared to young mice. In aged c-Jun OE/+ mice on the other hand, regeneration of DRG and motor neurons was restored to levels similar to those in young WT nerves.

In confirmation, counts of CGRP+ fibers in the sciatic nerve were performed 3 days after nerve crush, 3mm from the injury site (Fig.1 J). The number of fibers were reduced in aged WT nerves compared to young ones, but increased in the aged c-Jun OE/+ nerves.

These experiments confirm that the failure of repair cell function in older animals is accompanied by failure to fully upregulate c-Jun in Schwann cells after injury (Painter et al. 2014). Importantly, restoration of c-Jun elevation selectively in Schwann cells to that seen in young animals is sufficient to restore nerve regeneration to youthful levels.

### A MOUSE MODEL OF DISTAL STUMP DETERIORATION

We established a model of chronic denervation in mice, since previous studies have been carried out in rats. Sciatic nerves were cut followed by deflection of the proximal stump to leave the distal stump un-innervated for 1 week (short-term denervation) or 10 weeks (chronic denervation). mRNA levels for genes associated with denervated Schwann cells (*S100b, p75NTR, GDNF*, and *sonic hedgehog* (*Shh*)) declined substantially between 1 and 10 weeks of denervation (Fig.2 A). Western blots of p75NTR in the distal stump showed a rise to a maximum 1 week after injury and a decline thereafter to < 50% of peak levels at 10 weeks (Fig. 2 B) in line with that seen in rat (You et al. 1997).

**Figure 2:**
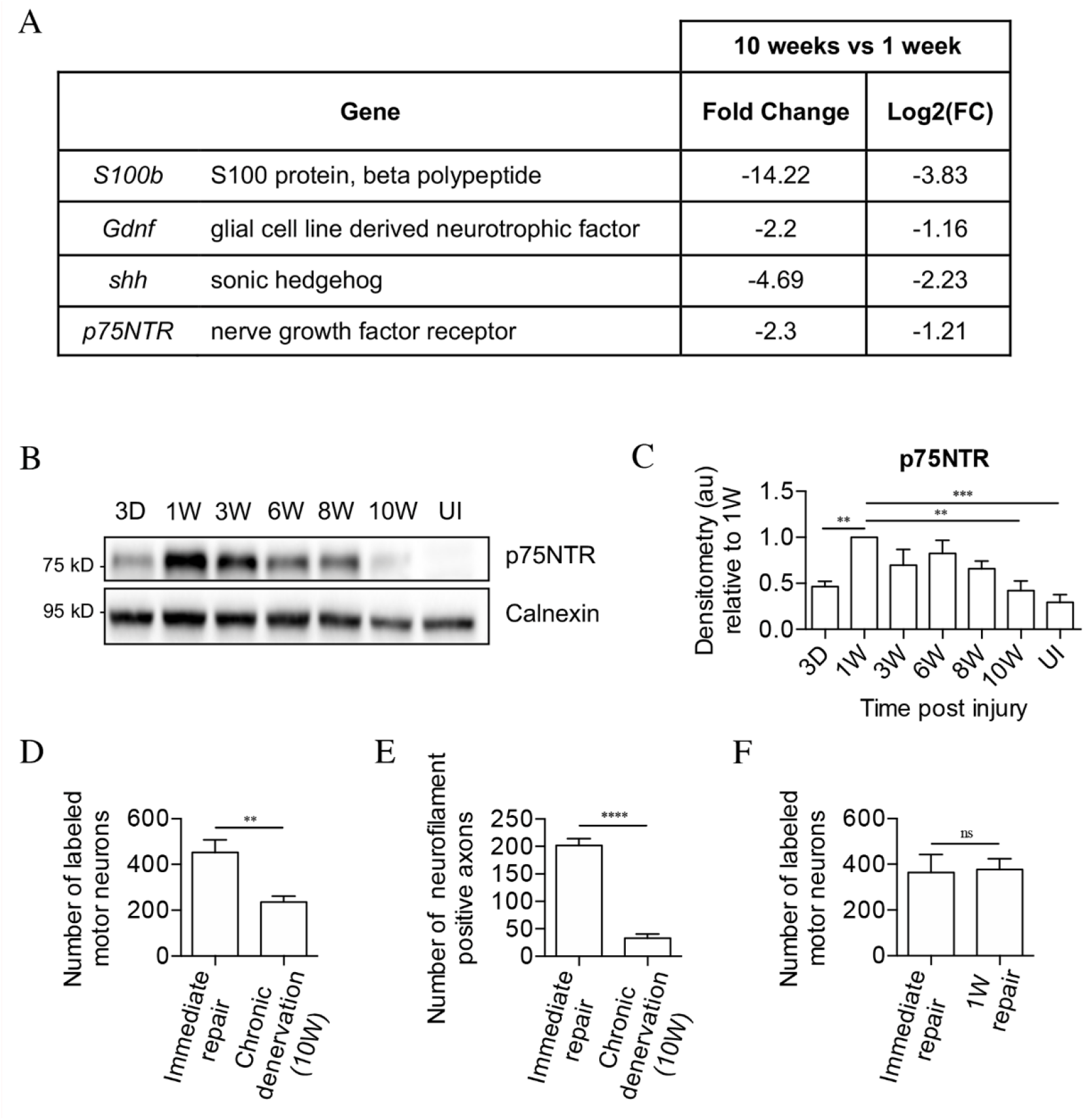
The mouse model of chronic denervation. (A) Analysis of RNA sequencing data showing decrease in gene expression during chronic denervation. (B) Representative Western blot showing p75NTR expression in uninjured (UI) nerves and distal nerve stumps following 3 days, and 1, 3, 6, 8 and 10 weeks of denervation. Quantification by densitometry is in (C). p75NTR peaks 1 week after injury and gradually declines during prolonged denervation. Data normalized to 1 week after injury. One-way ANOVA with Dunnett’s multiple comparison test; **p<0.005, ***p<0.001. n=4. (D) Counts of back-filled Fluorogold-labeled regenerating motor neurons following immediate repair or chronic 10 week denervation show a decrease in motor neuron regeneration into chronically denervated stumps. Unpaired Student’s t-test; **p= 0.0020. n=6 for each time point. (E) Counts of neurofilament+ axons mirrors the decline in regeneration observed with chronic denervation shown in *D*. Counts were performed on transverse sections taken 3mm from the repair site1 week after repair. Unpaired Stud ent’s t-test; ****p<0.0001. Immediate repair n= 5, chronic denervation n=4. (F) Counts of back-filled Fluorogold-labeled motor neurons showing similar numbers of regenerating neurons following immediate repair or repair after 1 week of denervation. Unpaired Student’s t-test; p=0.9. n=3. All numerical data represented as means ± SEM.

Regeneration through freshly cut and long-term denervated nerve stumps were compared by back-filling of spinal cord motor neurons. For this, acutely cut (immediate repair) or 10 week denervated tibial distal stumps were sutured to freshly cut peroneal nerves. Two weeks later, Fluorogold retrograde tracer was applied to the tibial stumps 4mm distal to the suturing site. One week after the application of tracer, the number of retrogradely labeled motor neurons in the spinal cord was counted. Only about half as many spinal cord motor neurons projected into the 10 week denervated stumps compared to the acutely transected stumps (Fig.2 D).

Failure of regeneration through 10 week denervated stumps was confirmed by immunohistochemistry and counting of neurofilament positive axons in similar experiments. Ten week denervated stumps contained many fewer regenerating neurofilament labeled axons than nerve stumps sutured immediately after transection (Fig.2 E).

Counting back-filled neurons following immediate repair or repair 1 week after transection, revealed similar number of regenerating neurons (Fig.2 F). This shows that the capacity of the distal stump to support regeneration declines between 1 and 10 weeks of denervation.

These experiments established a baseline for studying the effects of prolonged denervation on the capacity of mouse repair Schwann cells to support neuronal regeneration.

### c-JUN IS DOWN-REGULATED IN CHRONICALLY DENERVATED SCHWANN CELLS

We determined whether decline in Schwann cell c-Jun expression was involved in regeneration failure caused by chronic denervation, as seen during aging. Measuring c-Jun protein levels in distal stumps showed strong elevation 3 days and 1 week after sciatic nerve cut followed by a decline to ~40% of 1 week levels at 10 weeks (Fig.3 A, B). We verified that the c-Jun shown in these Western blots represented c-Jun in Schwann cells, using mice with conditional c-Jun inactivation selectively in Schwann cells (Arthur-Farraj et al. 2012) (Fig.3 C, D). Further, 1 and 10 week denervated stumps were compared using double immunofluorescent labelling with c-Jun antibodies and Sox10 antibodies to selectively identify Schwann cell nuclei. Ten week denervated stumps showed many more c-Jun negative, Sox10 positive nuclei, and the c-Jun labelling, where present, was also weaker (Fig.3 E).

**Figure 3:**
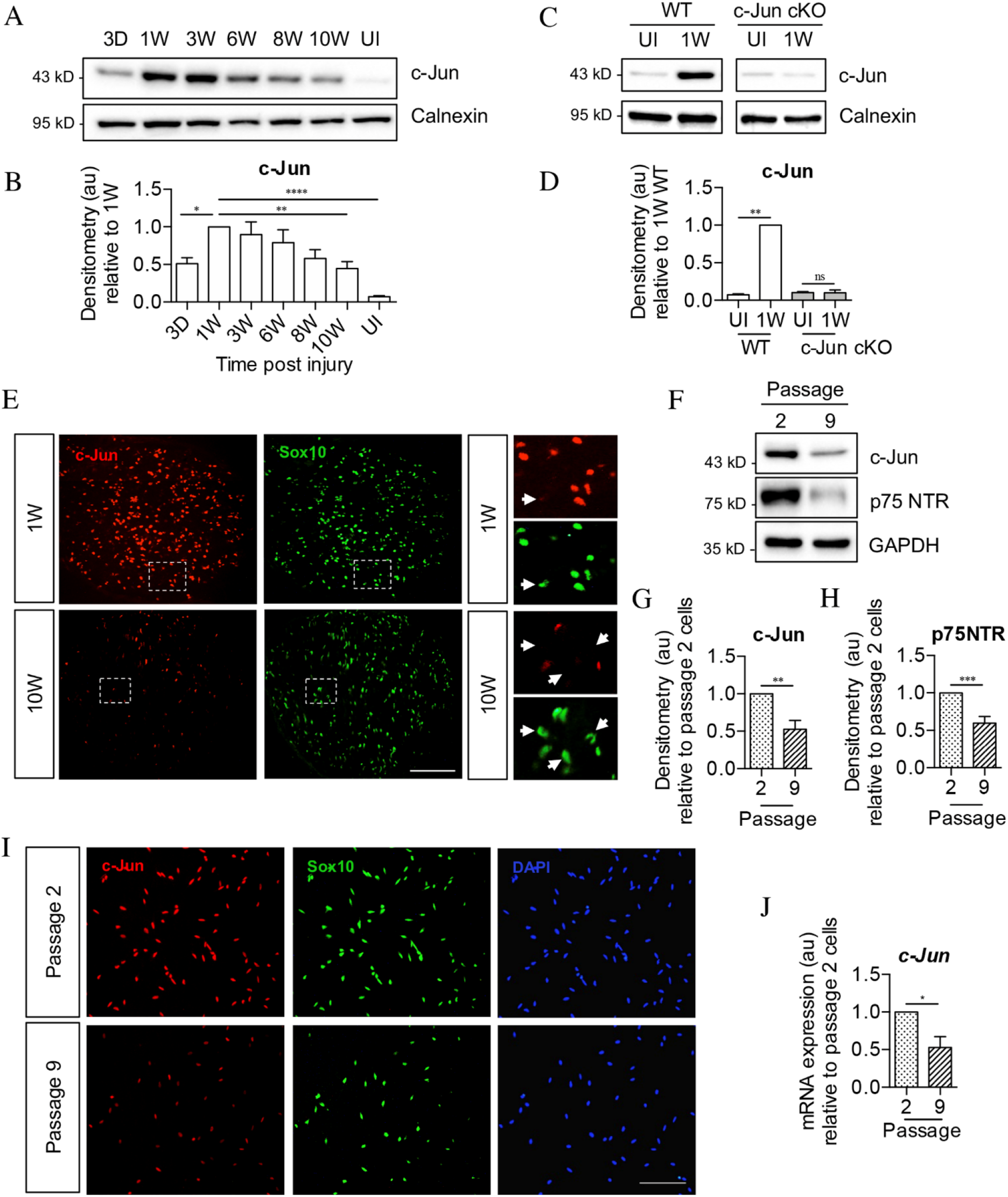
c-Jun declines in the distal nerve stump during chronic denervation and longterm culture. (A) Representative Western blot of c-Jun in WT uninjured (UI) nerves and distal stumps following 3 days and 1, 3, 6, 8 and 10 weeks of denervation. Quantification by densitometry is in (B), showing initial increase followed by a decline in c-Jun levels. Data normalized to 1 week post injury. One-way ANOVA with Dunnett’s multiple comparisons test; *p<0.05, **p<0.005, ****p<0.0001. n=5. (C) Representative Western blot comparing c-Jun expression 1 week after injury in WT and c-Jun cKO mice. Quantification is in (D), showing up-regulation of c-Jun in WT nerves but not in c-Jun cKO nerves, demonstrating that the c-Jun up-regulation after injury is Schwann cell specific. Data normalized to WT 1 week post injury. Two-way ANOVA with Sidak’s multiple comparison test; ****p<0.0001. n=5. (E) Representative immunofluorescence images of c-Jun/Sox10 double labelling in transverse sections of the distal stumps 1 and 10 weeks after cut. Boxed areas shown at higher magnification in right hand panels. Note loss of c-Jun protein from Schwann cell nuclei at 10 weeks (arrows). (F) Representative Western blot of c-Jun and p75NTR in Schwann cell cultures after 2 or 9 passages. Quantification is in (G and H), showing decline in c-Jun and p75NTR with time *in vitro*. Data normalised to passage 2. Unpaired Student’s t-test; **p=0.0023, ***p=0.0007. n=6 for c-Jun, n=7 for p75NTR. (I) c-Jun/Sox10 double labeling with nuclear marker DAPI after 2 or 9 passages. Note decline of nuclear c-Jun in passage 9 cells. (J) qPCR analysis showing reduction in c-Jun mRNA in Schwann cultures following 9 passages. Data normalized to passage 2. Unpaired Student’s t-test; *p=0.0299. n= 3. All numerical data represent means ± SEM.

The decline in c-Jun and p75NTR expression after long-term denervation *in vivo* was mimicked in Schwann cells deprived of axonal contact *in vitro*. Using purified Schwann cell cultures and Western blots, cells that had been maintained ~6 weeks *in vitro* (9 passages), contained less c-Jun and p75NTR protein compared to cells maintained ~10 days *in vitro* (2 passages) (Fig.3 F-H). The reduction in c-Jun in Schwann cell nuclei was confirmed using double immunofluorescent labelling with c-Jun and Sox 10 antibodies to identify Schwann cells (Fig.3 I). The levels of c-Jun mRNA also declined in long-term cultures (Fig.3 J) These *in vitro* experiments suggests that the decline in c-Jun and p75NTR during chronic denervation is not driven by endoneurial signals.

### C-JUN DOWN-REGULATION DURING CHRONIC DENERVATION IS PREVENTED IN C-JUN OE/+ MICE

The decline in c-Jun levels at the same time as repair cells lose capacity to support regeneration raises the questions of whether the functional deterioration of these cells is partly a consequence of c-Jun reduction, and whether repair cells, and regeneration through chronically denervated nerves, would be maintained if c-Jun reduction was prevented.

We addressed this using the c-Jun OE/+ mice examined earlier in experiments on aging (previous section; Fazal et al. 2017). The mice were 6-8 weeks old at the time of injury, corresponding to young mice in the study on aging. One and 3 week denervated distal stumps of c-Jun OE/+ and WT mice contained similar c-Jun protein levels (Fig.4 A, B). However, at 10 weeks, when c-Jun had declined in WT mice, c-Jun was maintained in c-Jun OE/+ mice at levels similar to those at 1 week. This was confirmed by double label c-Jun/Sox10 immunofluorescence (Fig.4 C, D). This showed similar c-Jun nuclear labeling in 1 week denervated stumps of WT and c-Jun OE/+ mice, while at 10 weeks, WT nerves showed reduced number of c-Jun positive nuclei and decreased labeling intensity. This decrease was prevented in c-Jun OE/+ nerves. (Fig.4 C, D).

**Figure 4:**
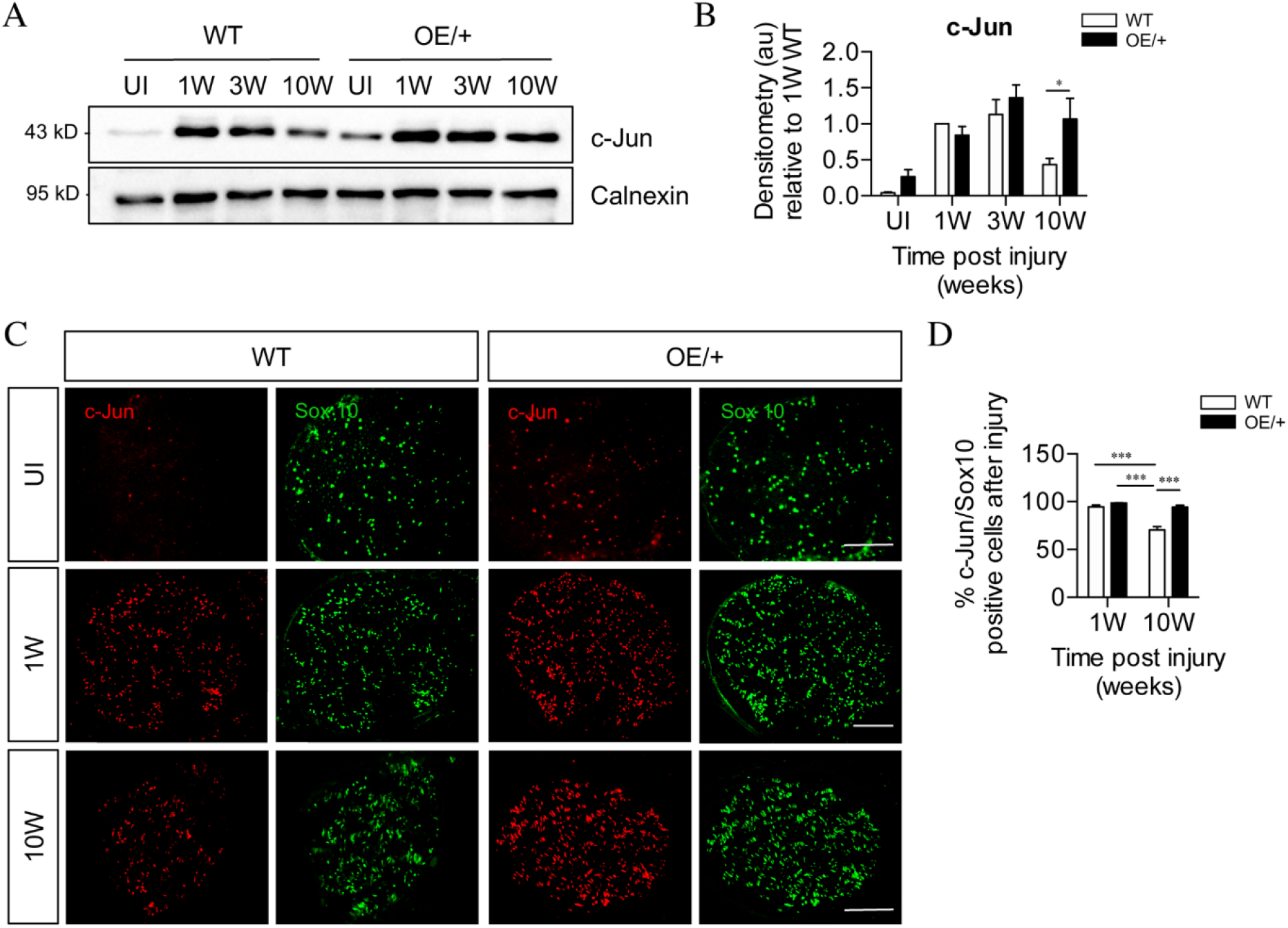
c-Jun expression is maintained in c-Jun OE/+ Schwann cells during chronic denervation. (A) Representative Western blots of c-Jun in WT and c-Jun OE/+ distal stumps after 1, 3 and 10 weeks of denervation. Quantification is in (B). In contrast to WT nerves, c-Jun OE/+ nerves maintain consistent levels of c-Jun during 10 week chronic denervation. Data normalized to WT 1 week post injury. Two-way ANOVA with Sidak’s multiple comparisons test; *p<0.05. n=5. (C) Representative images showing c-Jun/Sox10 double immunofluorescence in transverse sections of WT and OE/+ uninjured and injured distal stumps. Quantification by cell counting in (D). The c-Jun labeling of Sox10 positive nuclei in the two genotypes is comparable at 1 week, but reduced at 10 weeks in WT nerves only. Two-way ANOVA with Tukey’s multiple comparison test; *** P<0.001. n= 3. All numerical data represented as means ± SEM, all scale bars: 100μm.

These experiments indicate that in 1-3 week cut nerves, the maximum capacity of Schwann cells to express c-Jun protein is already reached in the WT, both genotypes showing a similar 80-100 fold elevation after injury. During chronic denervation, these high expression levels fall substantially in WT nerves, but not in c-Jun OE/+ nerves. The c-Jun OE/+ mice can therefore be used to test whether the regeneration failure induced by chronic denervation is due to the failure to maintain high c-Jun protein levels.

### MAINTAINING C-JUN LEVELS DURING CHRONIC DENERVATION PREVENTS REGENERATION FAILURE

Regeneration thorough chronically denervated distal stumps of WT and c-Jun OE/+ mice was compared using neuron back-filling. Acutely cut or 10 week denervated tibial nerves were sutured to acutely cut common peroneal nerves, and allowed to regenerate for 2 weeks prior to application of Fluorogold retrograde tracer. In WT mice, the number of DRG neurons projecting through 10 week cut nerves was only about half that projecting through acutely cut nerves. In 10 week cut c-Jun OE/+ nerves, however, this drop was not seen (Fig.5 A, B). Similar results were obtained for spinal cord motor neurons (Fig.5 C, D). Thus, both DRG and motor neurons regenerated as well through 10 week denervated c-Jun OE/+ nerves as they did through acutely cut WT nerves, suggesting that c-Jun OE/+ Schwann cells maintain their capacity to support regenerating neurons despite chronic denervation. In confirmation, regeneration into 10 week denervated distal stumps of the tibial nerve was examined with neurofilament staining, 1 week after the stumps were sutured to freshly cut peroneal nerves. Nearly 3 times more fibers were found in 10 week denervated c-Jun OE/+ nerves compared to 10 week denervated WT nerves (Fig.5 E).

**Figure 5:**
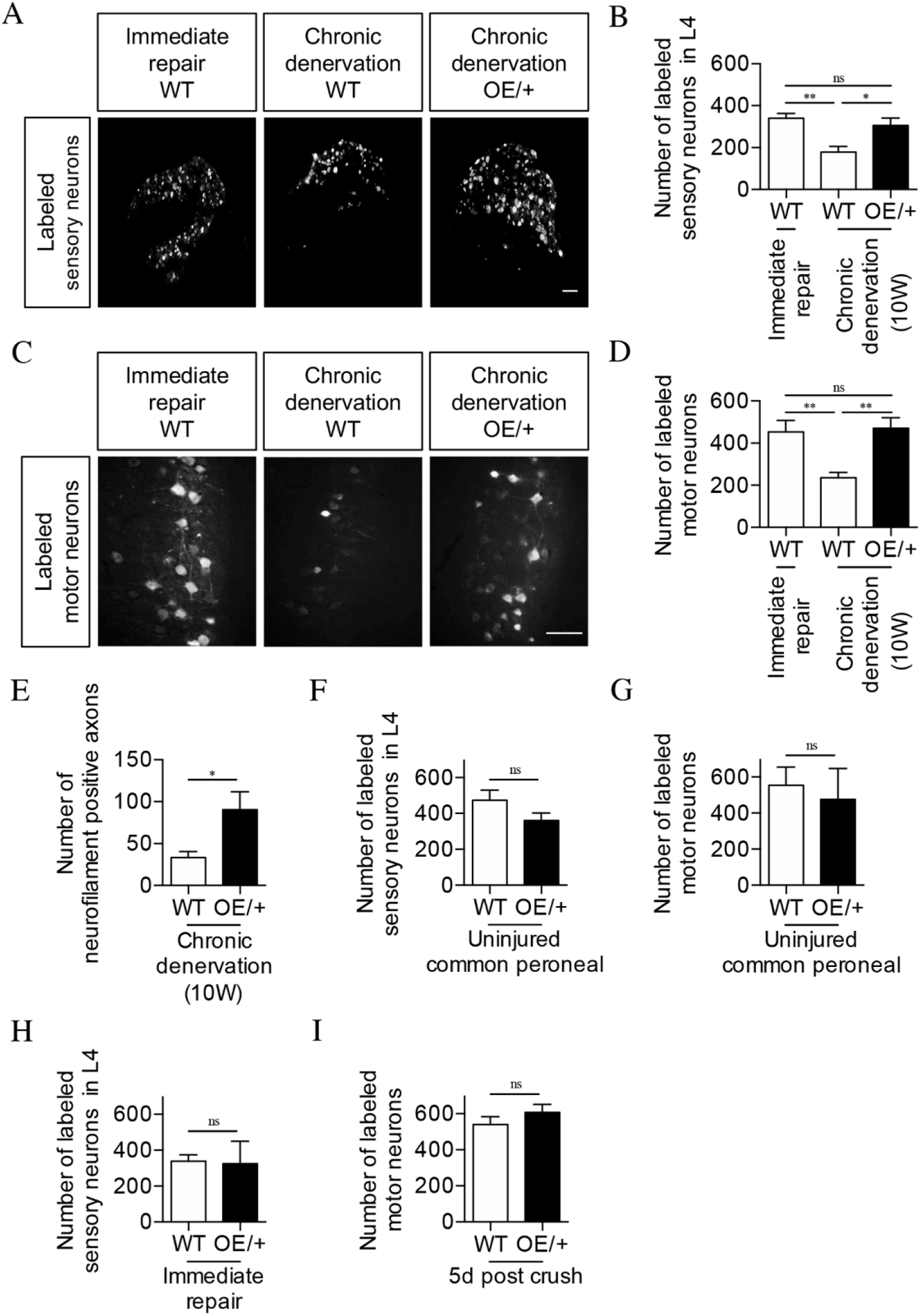
The regenerative capacity of c-Jun OE/+ nerves is maintained during chronic denervation. (A) Representative images showing Fluorogold-labeling of neurons in L4 DRGs of WT and c-Jun OE/+ mice after 2 weeks of regeneration into acutely transected (immediate repair) or chronically denervated distal stumps. Quantification by cell counting is in (B). The number of back-filled DRG neurons following regeneration through chronically denervated WT stumps was reduced, but maintained after regeneration through chronically denervated c-Jun OE/+ stumps. One-way ANOVA with Tukey’s multiple comparison test; **p<0.005, *p<0.05, ns non-significant. WT immediate repair and chronic denervation n= 6, OE/+ chronic denervation n=8. (C) Representative images showing Fluorogold-labeling of back-filled motor neurons in WT and c-Jun OE/+ mice after 2 weeks of regeneration into acutely transected (immediate repair) or chronically denervated distal stumps. Quantification is in (D), showing that compared to immediate repair, the number of labeled neurons is reduced after regeneration through chronically denervated WT stumps, but not after regeneration through chronically denervated c-Jun OE/+ stumps. One-way ANOVA with Tukey’s multiple comparison test; **p<0.005. WT immediate repair and chronic denervation n= 6, c-Jun OE/+ chronic denervation n=8. (E) Counts of neurofilament+ immunofluorescent axons in the distal stump 1 week after repair following chronic denervation of WT and c-Jun OE/+ nerves. Unpaired Student’s t-test; *p= 0.0334. WT n=4, c-Jun, OE/+ n=3. (F, G) Counts of Fluorogold-labeled sensory (F) and motor (G) neurons in WT and c-Jun OE/+ mice following transection with immediate application of tracer. The number of back-filled sensory (p=0.1872) and motor (p=0.7153) neurons is similar. Unpaired Student’s t-tests, n=3 for each experimental condition. (H) The number of back-filled sensory neurons in WT and c-Jun OE/+ mice is similar after transection followed by immediate repair. Unpaired Student’s t-test; p= 0.9195. n=3. (I) The number of labeled motor neurons in WT and c-Jun OE/+ mice is similar when tracer was applied 5 days after sciatic nerve crush. Unpaired Student’s t-test; p=0.312. WT n=5, OE/+ n=4. All numerical data represented as means ± SEM, all scale bars: 100μm.

We verified that the back-filling paradigm worked as expected in c-Jun OE/+ mice. First, peroneal nerves in WT and c-Jun OE/+ mice were transected followed by immediate application of tracer to the injured proximal stump. The number of back-filled DRG and motor neurons in c-Jun OE/+ mice was similar to that in WT animals (Fig.5 F, G). This does not measure regeneration, but indicates that a comparable number of DRG and motor neurons project through the normal uninjured peroneal nerve in the two mouse lines. Second, although c-Jun levels in WT and c-Jun OE/+ mice diverge after 10 weeks of denervation, they are high, and similar, early after injury, when the capacity of WT nerves to support regeneration is optimal. In line with this, the regeneration support provided by WT and c-Jun OE/+ nerves was similar early after injury. Thus, in back-filling experiments no significant difference was seen between WT and c-Jun OE/+ mice in the numbers of DRG or motor neurones that regenerated through acutely transected distal stumps (immediate repair) when the tracer was applied 2 weeks after transection/repair. (Fig.5 H and not shown). Similarly, comparable numbers of back-filled motor neurons were obtained in WT and c-Jun OE/+ mice when back-filling was used to quantify regeneration 5 days after sciatic nerve crush (Fig.5 I).

Together, these experiments indicate that long-term absence of axonal contact leads to reduction in Schwann cell c-Jun, and deterioration of repair cell function, resulting in regeneration failure, a failure that can be corrected by restoring c-Jun levels.

### IMPROVED REGENERATION IN C-JUN OE+/- MICE IS UNLIKELY TO RELATE TO ALTERED CELL NUMBERS

To test whether the differential regeneration rates could be due to altered cell numbers, Schwann cell, macrophage and fibroblast nuclei were counted in tibial nerves by electron microscopy.

In experiments comparing young and aging animals, cells in the distal stump were counted 4 days after nerve cut without regeneration. Schwann cell and macrophage numbers and density were similar irrespective of genotype or age (Fig.6 A-C). There was some decrease in fibroblast density in aged mice, but density and numbers were similar irrespective of genotype (Fig.6 D, E). In aged mice, transverse profiles of the tibial nerve were larger because of increased endoneurial connective tissue, but no difference was seen between the genotypes (Fig.6 F).

**Figure 6:**
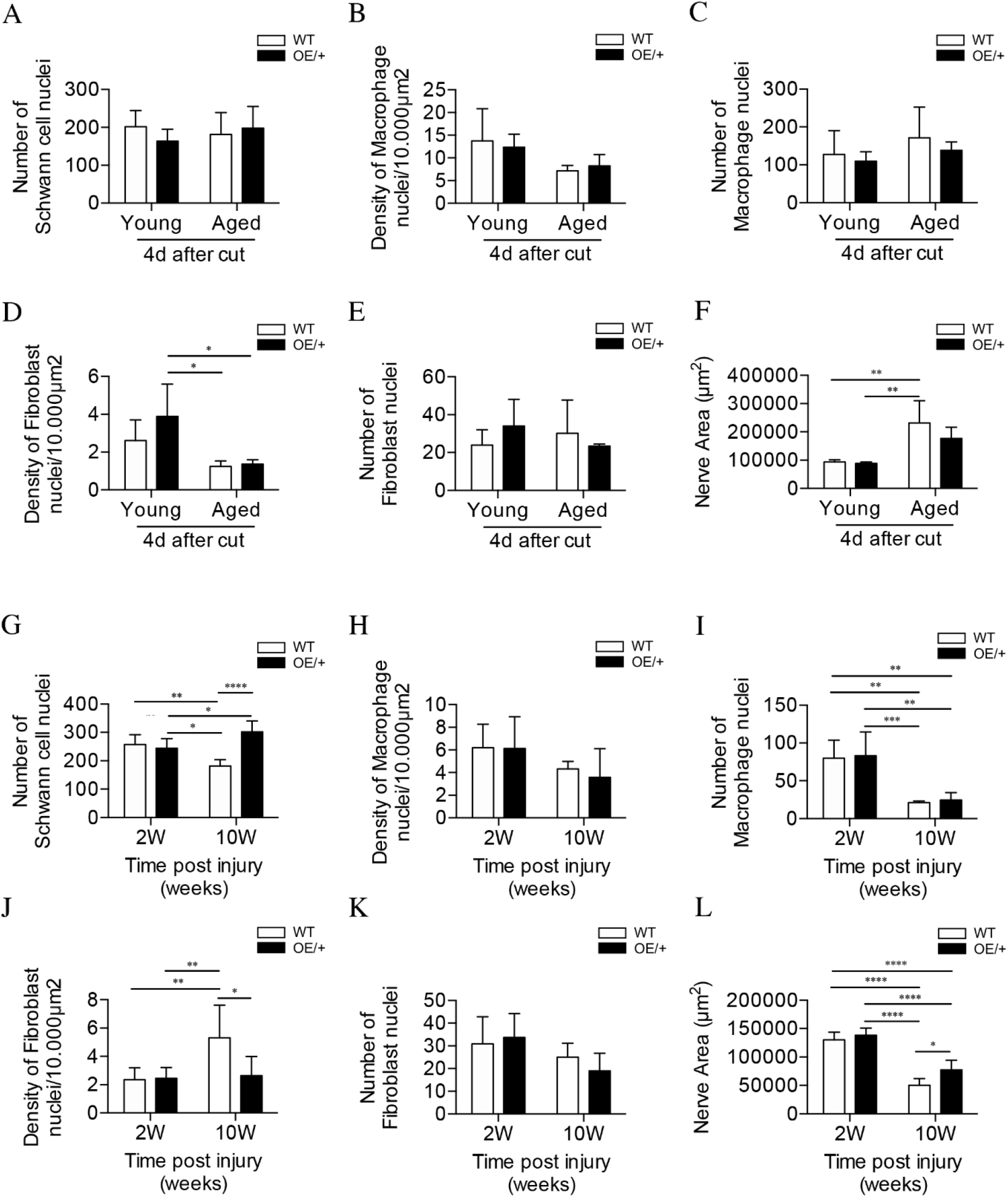
Cell number and nerve size in injured WT and c-Jun OE/+ nerves. Cell nuclei were counted in whole transverse profiles of the tibial nerve, 5mm from the injury site, using the electron microscope. (A) Schwann cell numbers in young and aged WT and c-Jun OE/+ nerves. (B) Macrophage density and (C) number in young and aged WT and c-Jun OE/+ nerves. (D) Fibroblast density and (E) number in young and aged WT and c-Jun OE/+ nerves: *p<0.05. (F) Whole transverse profiles of the tibial nerve were measured for the nerve area of young and aged WT and c-Jun OE/+ nerves: **p<0.005. For counts in A-F, n=4 for each condition. (G) Schwann cell numbers in 2 and 10 week cut nerves of WT and c-Jun OE/+ mice; *p<0.05, **p<0.005, ****p<0.0001. (H) Macrophage density and (I) number in 2 and 10 week cut nerves of WT and c-Jun OE/+ mice: **p<0.005, ***p<0.001. (J) Fibroblast density and (K) number in 2 and 10 week cut nerves of WT and c-Jun OE/+ mice: *p<0.05, **p<0.005. (L) Whole transverse profiles of the tibial nerve were measured for the nerve area of 2 and 10 week cut nerves of WT and c-Jun OE/+ mice: *p<0.05, ****p<0.0001. For counts in G-L, 2 week WT and c-Jun OE/+ n=7, 10 week WT and c-Jun OE/+ n=5. All number data represented as means ± SEM analysed by two-way ANOVA with Tukey’s multiple comparison test.

In the experiments on chronic denervation, cells were counted in the distal stumps 2 and 10 weeks after cut without regeneration 5 mm from the injury site. There were ~250 Schwann cell nuclei in 2 week stumps, representing ~2.5 fold increase from uninjured nerves (Fazal et al. 2017) (Fig.6 G). In the WT, Schwann cell numbers declined by ~30% during chronic denervation. This drop was prevented in c-Jun OE/+ nerves (Fig.6 G). The density of macrophages was similar irrespective of genotype or length of denervation (Fig.6 H), with numbers declining, irrespective of genotype, during chronic denervation (Fig.6 I). Two weeks after injury, fibroblast density in WT and c-Jun OE/+ mice was similar. Fibroblast density was elevated in 10 week stumps of WT, but not c-Jun OE/+, mice (Fig.6 J). Fibroblast numbers remained unchanged between genotypes following denervation (Fig.6 K). Chronically denervated stumps of both genotypes had reduced nerve area, although this difference was less evident in c-Jun OE/+ nerves (Fig.6 L).

These counts indicate that altered cell numbers are not a significant reason for improved regeneration in aged c-Jun OE/+ nerves. In chronically denervated nerves, the normal loss of Schwann cells in WT nerves is prevented in c-Jun OE/+ nerves. Even in 10 week WT stumps, however, cell numbers remain well above those in uninjured nerves (see Discussion).

### ANALYSIS OF GENE EXPRESSION IN DISTAL NERVE STUMPS OF WT AND C-JUN OE/+ MICE

#### Young and aging mice

RNA sequence analysis was performed on uninjured and 3 day cut sciatic nerves of young (6-8 weeks) and aged (11-12 months) WT mice, and aged c-Jun OE/+ mice. Global gene expression was analysed comparing (i) uninjured nerves, (ii) 3d cut nerves, and (iii) the injury response (3d cut vs uninjured) in young and aged mice (Fig.7 - Figure Supplement 1 A).

In uninjured WT nerves, out of 15,995 genes present, 1,477 genes were differentially expressed between young and aged mice. Of these, 1,154 were up-regulated and 323 down-regulated in aging mice compared to young ones (FC >2; and FDR 0.05) (Fig. 7 - Figure Supplement 2 A; Supplementary Table 1 A). We tested whether the 173 genes we previously identified as c-Jun regulated injury genes (Arthur-Farraj et al. 2012) were implicated in these age-dependent differences. In the present data set, 138 of the 173 genes were present (Supplementary Table 2). They showed a strong enrichment among the 1477 genes differentially expressed between young and aging mice (Fig.7 A).

**Figure 7:**
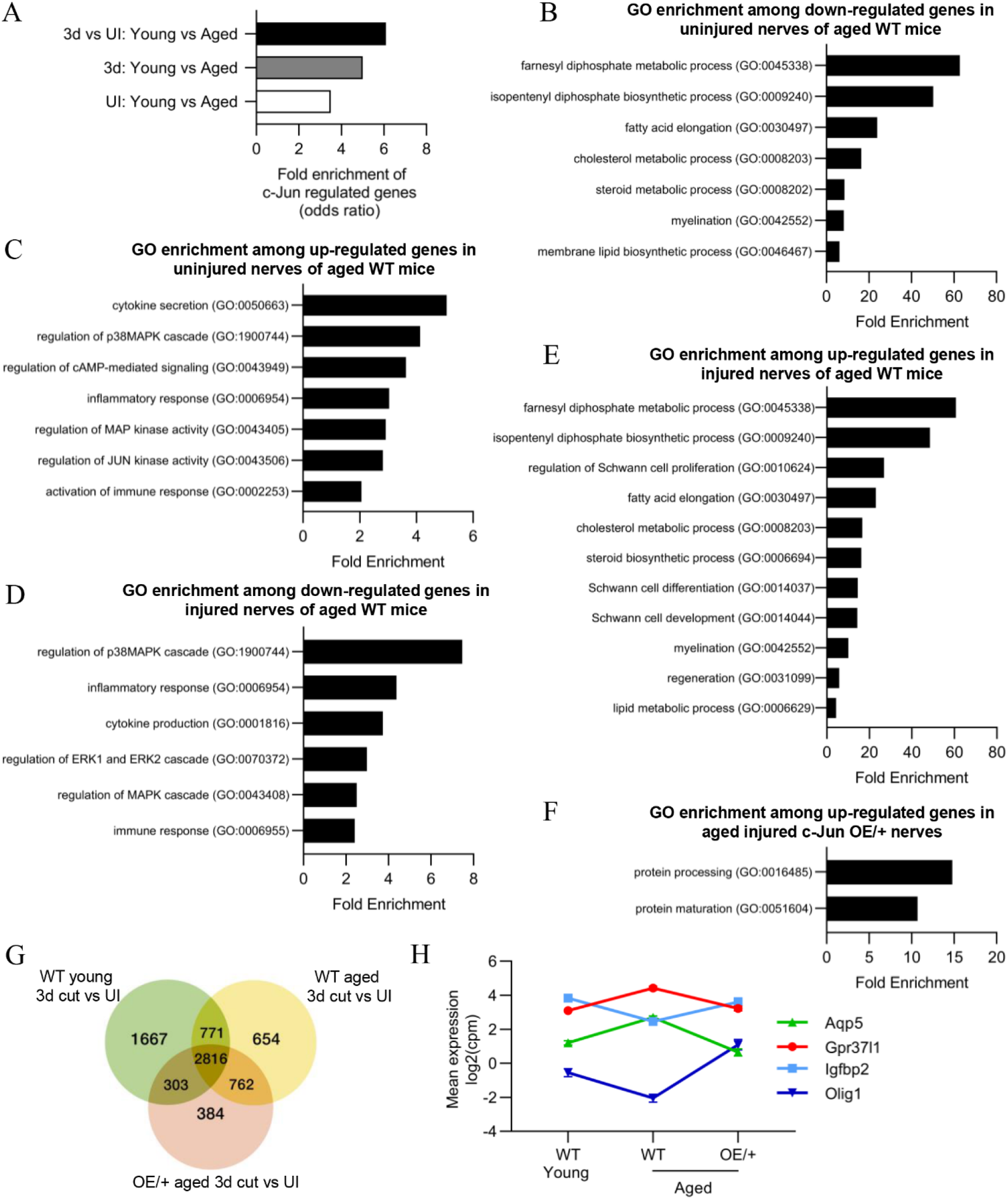
Bioinformatics analysis of RNA seq. data from young and aged nerves. (A) Over-representation analysis showing enrichment of c-Jun regulated genes in various WT injury paradigms. p= 3.2×10^-8^ for UI young vs aged; p=1×10x ^-7^ for 3 day cut young vs aged; p=2.3×10^-13^ for the injury response. p-values computed by one-sided Fisher’s exact test. (B) GO terms down-regulated and (C) up-regulated in uninjured nerves of aged WT mice. (Absolute fold change >2 and FDR<0.05). (D) GO terms down-regulated and (E) up-regulated in the injury response of aged WT mice. (Absolute fold change >2 and FDR<0.05). (F) When aged c-Jun OE/+ and WT nerves are compared, genes associated with protein processing (FDR = 0.00318) and maturation (FDR = 0.0153) are significantly enriched in aged c-Jun OE/+ nerves compared to aged WT. (G) Venn diagram showing numbers of differentially expressed genes between young and aged 3 day cut WT nerves and aged 3 day cut OE/+ nerves, compared to their uninjured counterparts. (H) Mean expression of 4 c-Jun regulated genes with significantly different expression between young and aged WT nerves but not between young WT and aged c-Jun OE/+ nerves. (Absolute fold change >2 and FDR<0.05).

In 3d cut WT nerves, of 17,334 genes present, 398 genes were differentially expressed between young and aging mice. Of these, 268 were up-regulated and 130 down-regulated in aging nerves (Fig.7 - Figure Supplement 2 B; Supplementary Table 1 B). This gene set contains candidate genes responsible for the difference in regeneration support provided by young and aged Schwann cells (Painter et al. 2014 and present experiments). In agreement with Painter et al. (2014), trophic factors such as GDNF, BDNF, NGF, erythropoietin and FGF were not among the differentially expressed genes. This suggests that expression of trophic factors often implicated in regeneration may not explain different regeneration between young and aged mice. The 138 c-Jun regulated injury genes (Arthur-Farraj et al. 2012) were highly enriched among the age-regulated genes (Fig.7 A).

Examining the injury response (3 d cut vs uninjured nerve), 822 genes showed significant difference when young and aging WT mice were compared (Fig.7 – Figure Supplement 2 C; Supplementary Table 1 C). The 138 c-Jun regulated injury genes were strongly enriched among these 822 genes (Fig.7 A).

Gene set enrichment analysis (GSEA) on the above conditions showed that the 138 c-Jun genes were highly enriched among the genes down-regulated in aging uninjured nerves and in aging 3 day cut nerves (Fig. 7 - Figure Supplement 1 B, C). When the injury response (3d cut vs uninjured) of young and aging WT mice was analysed, the strongest enrichment of c-Jun genes was observed in genes upregulated in young nerves (Fig. 7 - Figure Supplement 1 D). While c-Jun genes are also upregulated in aged nerves post-injury, their enrichment was not as high as in young nerves.

These correlations between enrichment of c-Jun regulated genes and Schwann cell age suggest that the c-Jun regulated repair program is disproportionately vulnerable during the aging process.

Gene ontology (GO) analysis showed that in aged uninjured WT nerves, down-regulated genes were largely involved in lipid metabolism, as well as myelination, while genes involved in the immune system were prominent among those up-regulated (Fig.7 B, C; reviewed in Melcangi et al. 1998; 2000; Büttner et al. 2018). Similar analysis of the injury response (3 day cut vs uninjured) showed reduced activation of immune genes in aged WT nerves (Scheib and Höke 2016; Büttner et al. 2018). In aged nerves, MAPK pathways were also suppressed while lipid metabolism and Schwann cell differentiation genes were enhanced Fig.7 D, E). Together this indicates suppressed Schwann cell reprogramming and repair cell activation in nerves of older WT mice. Testing the effects of enhanced c-Jun expression on the aged injury response, we found that pathways associated with protein processing and maturation were up-regulated in aged c-Jun OE/+ nerves compared with aged WT nerves (Fig.7 F).

To further determine genes that may contribute to the restoration of regeneration in aged c-Jun OE/+ mice, the injury responses in young and aged WT mice and aged c-Jun OE/+ mice were compared (Fig.7 G). Of particular interest are the 303 genes that show significant injury response in young WT mice but not in aging WT mice, but are again significantly regulated by restoring c-Jun to youthful levels in aging c-Jun OE/+ mice (Supplementary Table 1 D).

Among the 138 c-Jun regulated genes, we looked for a correlation between a failure and restoration of gene expression on the one hand, and failure and restoration of regeneration on the other. In 3 day cut aged WT nerves, where regeneration fails, 16 c-Jun regulated genes were differentially expressed compared to 3 day cut young WT nerves. Four of these, Aqp5, Gpr37L1, Igfbp2, and Olig1, were restored in aged c-Jun OE/+ nerves, where regeneration is restored (Fig 7 H). Thus, in aging mice, both regeneration failure and the expression defect of these four genes was restored to levels in young mice, by elevating c-Jun levels.

#### Chronic denervation

Gene expression was examined in uninjured nerves and in 1 and 10 week cut sciatic nerves of WT and c-Jun OE/+ mice (Fig. 8 - Figure Supplement 1 A). Expression of 1581 genes changed significantly during chronic denervation; Figure 8 - Figure Supplement 2; Supplementary Table 1 E). In 10 week cut nerves, 601 of these genes were down-regulated, including genes associated with repair cells such as GDNF, Shh and p75NTR, while 980 genes were up-regulated. The 138 c-Jun regulated genes showed a highly significant 5.8 fold enrichment (p=2.2×10^-16^) among the 1581 genes regulated during chronic denervation. GSEA enrichment analysis showed that c-Jun genes were some of the most down-regulated genes during chronic denervation. (Fig.8 - Figure Supplement 1 B).

GO analysis showed that the major genes down-regulated during chronic denervation in WT nerves involved the cell cycle, DNA replication and repair. Glial cell differentiation genes and MAPK pathways, potential activators of c-Jun, were also suppressed (Fig.8 A). Chronic denervation involved a prominent up-regulation of neuro-glia signaling genes (chiefly related to GABA and adrenergic signalling), but also negative regulators of differentiation, Notch and cAMP signalling (Fig.8 B). To test the effects of maintaining c-Jun protein levels during the 10 week chronic denervation, we identified genes differentially expressed between 10 week cut WT and c-Jun OE/+ nerves (Fig.8 C). This showed strong up-regulation of pathways involved in PNS and Schwann cell development and differentiation in c-Jun OE/+ mice.

**Figure 8:**
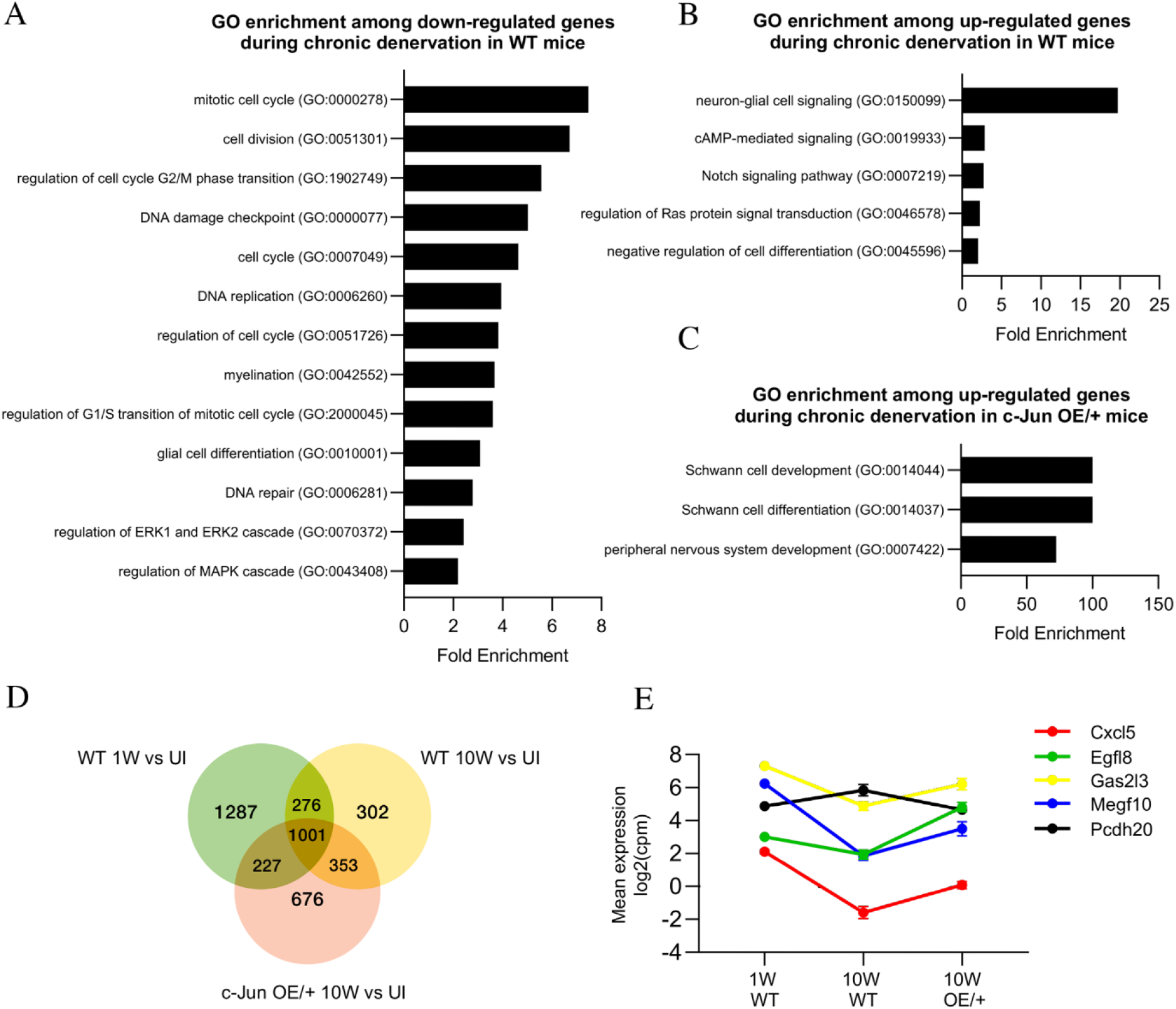
Bioinformatics analysis of RNA seq. data from acutely and chronically denervated nerves. (A) GO terms down-regulated and (B) up-regulated in WT nerves during chronic denervation(C) When chronically denervated c-Jun OE/+ and WT nerves were compared, GO terms associated with Schwann cell differentiation (FDR = 0.00397) and PNS development (FDR = 0.0173) were enriched in c-Jun OE/+ nerves. (D) Venn diagram showing numbers of differentially expressed genes between WT nerves following acute (1 week) and chronic (10 week) denervation and c-Jun OE/+ nerves following chronic denervation, compared to their uninjured counterparts. (E) Mean expression of 5 c-Jun regulated genes with significantly different expression between acute and chronic WT nerves, but not between acute WT and chronic c-Jun OE/+ nerves. (Absolute fold change >2 and FDR<0.05).

To further determine genes that may contribute to the restoration of regeneration in aged c-Jun OE/+ mice, the injury response in the three situations analysed in the regeneration experiments, WT 1 week, WT 10 weeks and c-Jun OE/+ 10 weeks, was compared (Fig.8 D). A point of interest are the 227 genes that showed an injury response in WT 1 week nerves and in c-Jun OE/+ 10 week nerves, both of which show fast regeneration, but no injury response in WT 10 week nerves, where regeneration is slow (Supplementary Table 1 F).

As when studying aged mice, we looked among the 138 c-Jun regulated injury genes for candidates involved in decreased regeneration in 10 week cut WT nerves and the restoration of regeneration in 10 week cut c-Jun OE/+ nerves. Fifty of the 138 genes changed expression during chronic denervation in WT nerves, where regeneration is poor. In chronically denervated c-Jun OE/+ nerves, where regeneration is restored, expression levels were restored, completely or partially, in the case of five of these genes, Cxcl5, Egfl8, Gas213, Megf10 and Pcdh20 (Fig.8 E). These correlations provide a basis for considering these genes as candidates down-stream of c-Jun for involvement in the restoration of regeneration in chronically denervated nerves of c-Jun OE/+ mice.

**Figure 7 - Figure Supplement 1.:**
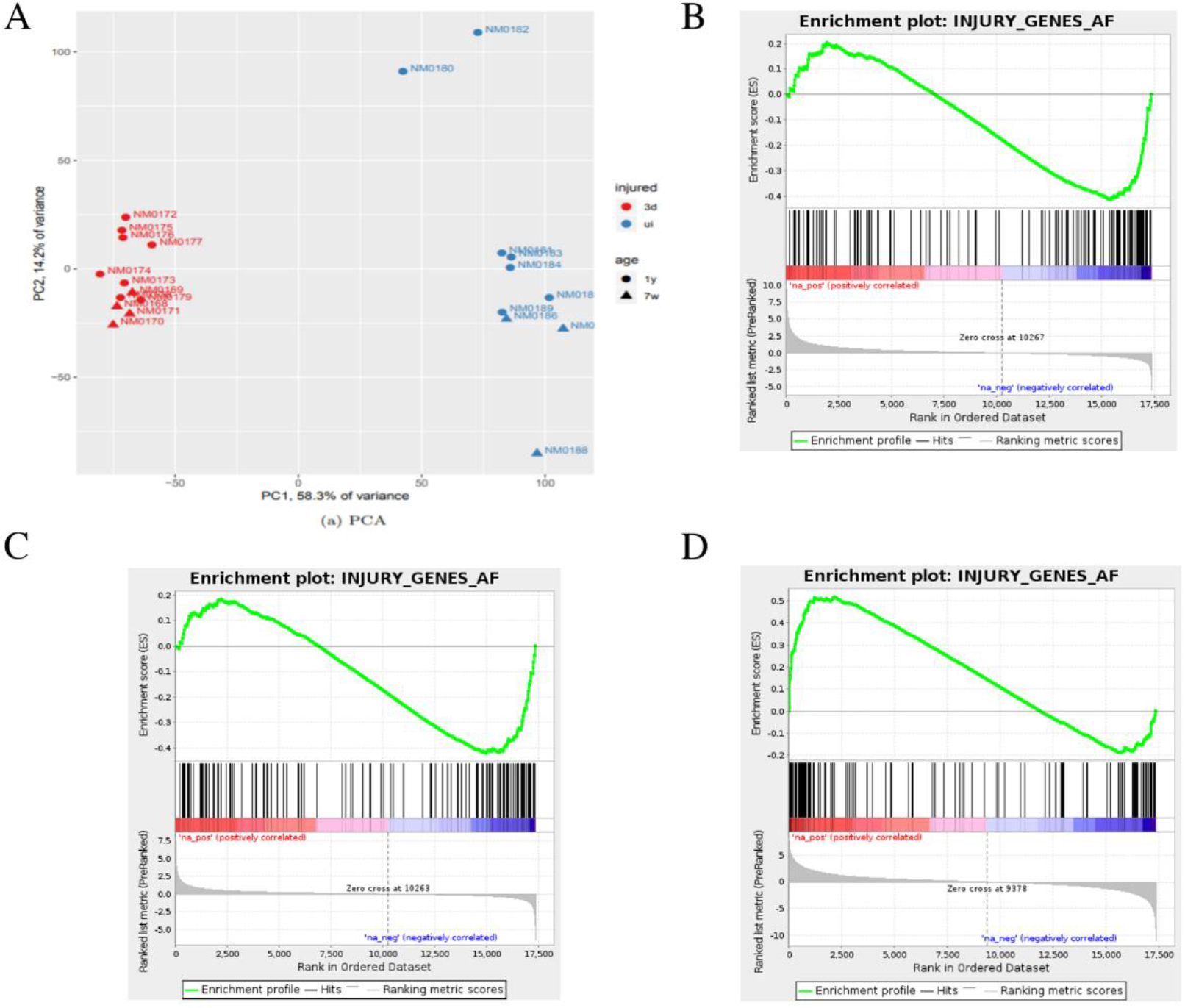
Bioinformatics analysis of aged and young nerves following injury. (A) Principal component analysis (PCA) shows that the key source of variance in our aging analysis is injury status with samples clustering together after injury (red) regardless of age or genotype. (B) Gene set enrichment analysis (GSEA) of uninjured young and aged WT nerves found enrichment in the down-regulation of c-Jun regulated genes in aged nerves. Normalized enrichment score (NES) = −1.85, q-value = 0.0. (C) GSEA analysis comparing injured old and young nerves found a similar enrichment of c-Jun genes in the down-regulation of these genes, suggesting a failure of c-Jun regulation in aged nerves. NES= −1.61, q-value = 0.0. (D) GSEA of young and aged nerves compared to their uninjured controls and then to each other to examine the differential expression between the two in response to injury. Analysis found that c-Jun gene up-regulation was highly enriched by injury. NES= 2.01, q-value = 0.0.

**Figure 7 - Figure Supplement 2.**
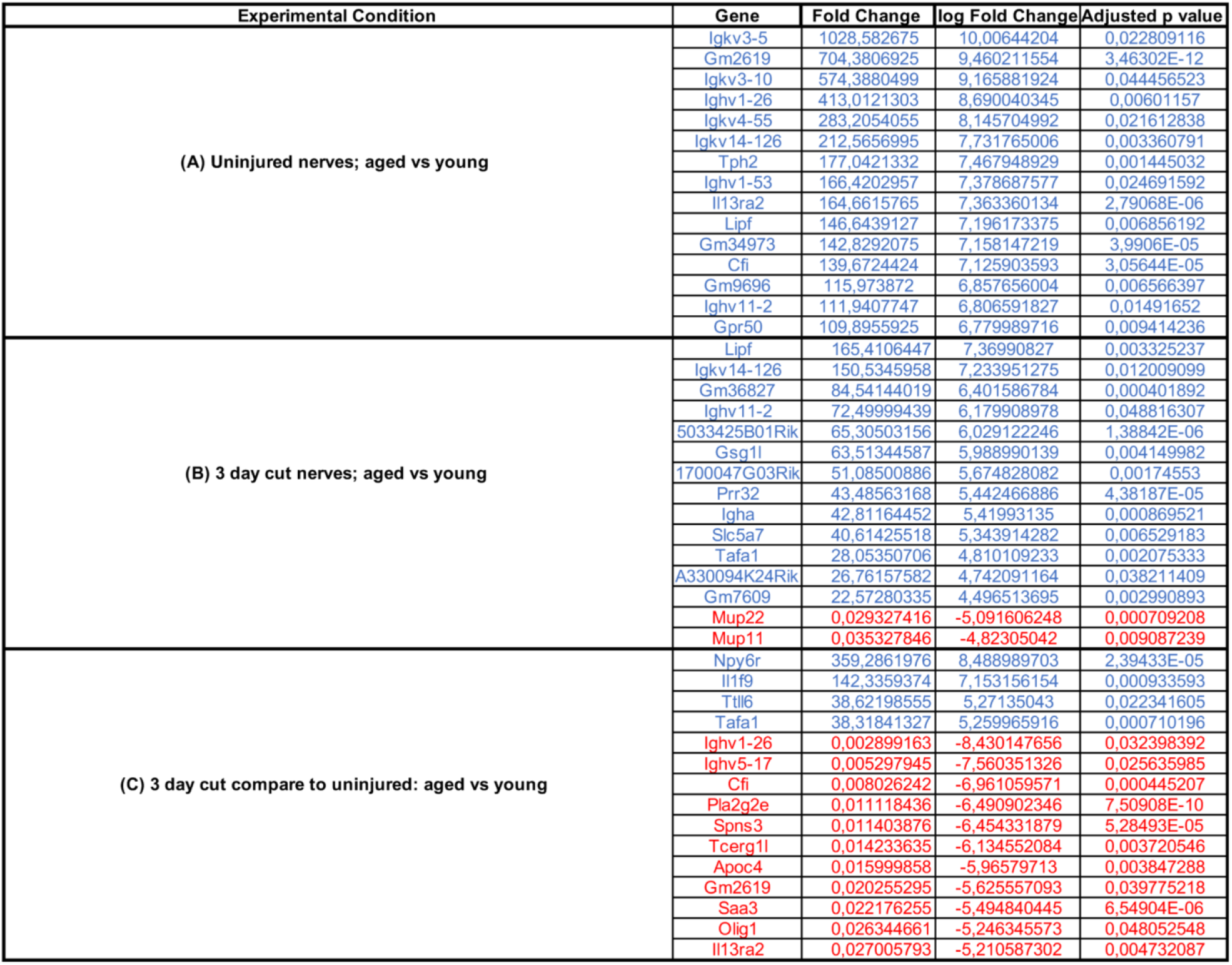
The 15 most regulated genes in the tibial nerve of WT mice during aging. (A, B) Genes expressed at higher levels in aged mice under the conditions indicted are in blue (top), while genes with reduced expression in aged mice are in red (bottom). (C) Genes that respond more strongly to injury in aged mice are in blue (top), while genes with weaker injury response are in red (bottom).

**Figure 8 - Figure Supplement 1.**
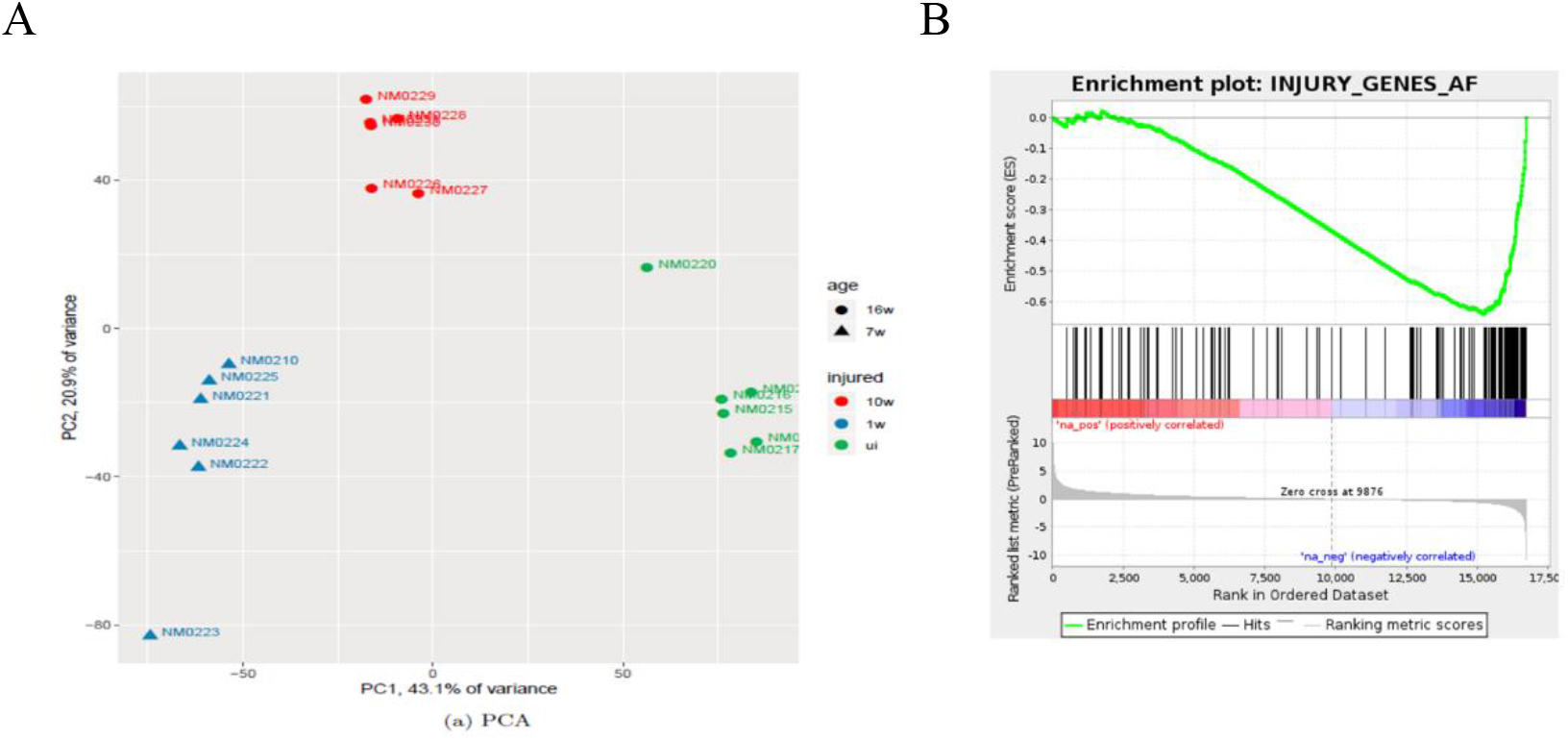
Bioinformatics analysis of short-term and chronically injured nerves. **(A)** PCA showing the key sources of variance in our chronic injury analysis are not only injury but time post injury with samples clustering based on length of denervation regardless of genotype. **(B)** GSEA of WT chronically denervated nerves found that c-Jun genes were some of the most down-regulated genes during chronic denervation NES = −2.5, q-value = 0.0.

**Figure 8 - Figure Supplement 2.**
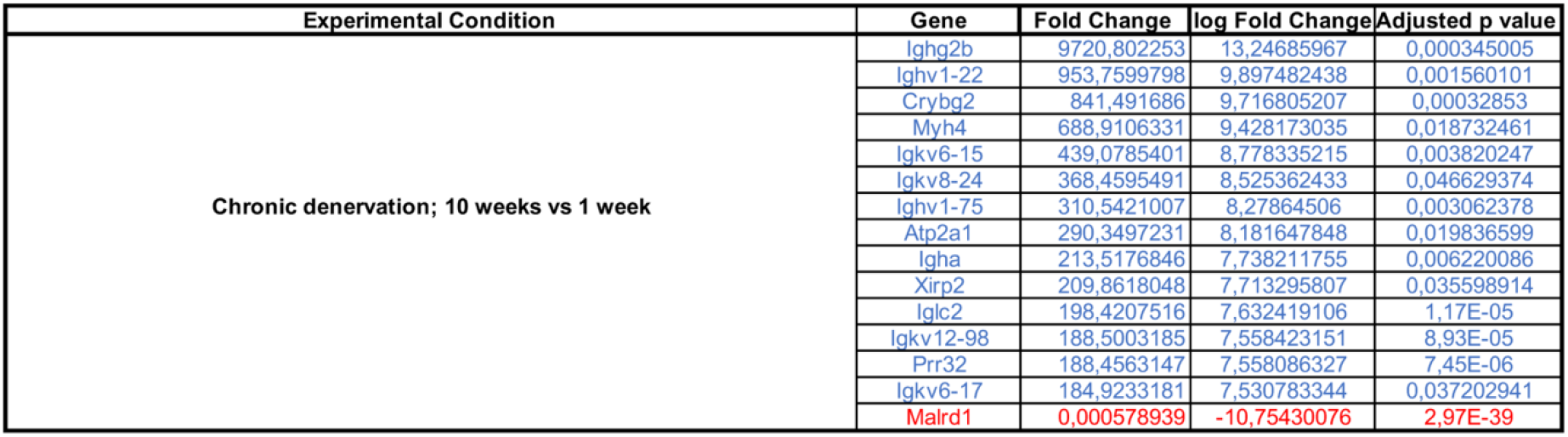
The 15 most regulated genes in the tibial nerve during chronic denervation. Genes expressed at higher levels after chronic denervation are in blue (top), while genes with reduced expression are in red (bottom).

### SHH SIGNALING SUPPORTS C-JUN EXPRESSION

Seeking mechanisms that promote c-Jun expression in denervated Schwann cells, we considered signaling by Shh, a gene that is not expressed in developing Schwann cells or in intact nerves, but strongly up-regulated in repair Schwann cells after injury (Lu et al. 2000; Zhou et al. 2000; Arthur-Farraj et al. 2012; Hsin-Pin et al. 2015). First, we analysed Shh cKO mice, in which Shh is electively inactivated in Schwann cells and in which nerve ultrastructure appears normal as expected (J Svaren unpublished). We found that in the mutants, c-Jun protein and phosphorylated c-Jun were decreased in the distal stump of 7 day transected sciatic nerve, (Fig.9 A-D). Levels of p75NTR protein, which is positively regulated by c-Jun in Schwann cells (Arthur-Farraj et al. 2012), were also reduced 7 days after injury in the mutants (Fig.9 E, F). Substantiating these observations, two Shh signaling agonists, SAG and purmorphamine, upregulated c-Jun protein in purified Schwann cell cultures (Fig.9 G-J). This was confirmed using c-Jun/Sox10 double immunolabeling (Fig.9 K). SAG also increased the expression of two c-Jun target genes that promote nerve regeneration, GDNF and BDNF (Fig.9 L, M). Further, SAG promoted another effect of c-Jun, that of enhancing the elongated bi-or tri-polar shape in vitro that reflects the elongation and branching of repair cells in vivo (Arthur-Farraj et al. 2012; Gomez-Sanchez et al. 2017) (Fig.9 N).

**Figure 9:**
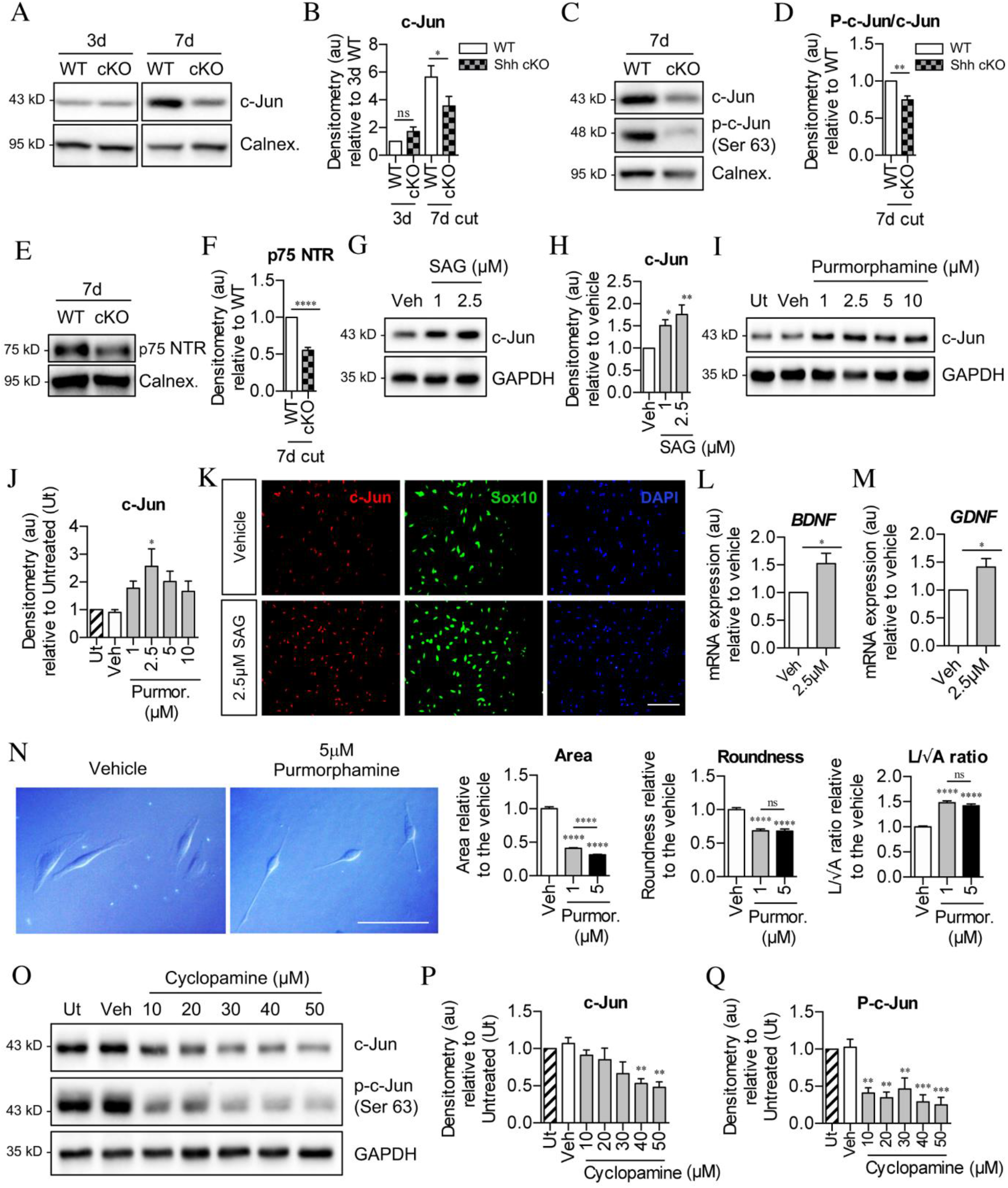
Sonic hedgehog plays a role in c-Jun activation in Schwann cells via autocrine signaling. (A) Representative Western blot showing c-Jun expression in WT and Shh cKO (cKO) nerves 3 and 7 days after cut. Densitometry shown in (B). Data normalized to WT 3 days post cut. Two-way ANOVA with Sidak’s test; *p<0.05. n=5 for each genotype. (C) Representative Western blot showing c-Jun and phosphorylated c-Jun (p-c-Jun in WT and Shh cKO distal nerve stumps 7 days post cut. Densitometry shown in (D). Data normalized to WT 7 days post injury. Unpaired Student’s t-test; **p=0.0014, n=5 for each genotype. (E) Representative Western blot showing p75NTR protein in WT and Shh cKO nerves 7 days post cut. Densitometry shown in (F). Data normalized to WT 7 days post injury. Unpaired Student’s t-test; ****p=<0.0001. n=5 for each genotype. (G) Representative Western blot showing c-Jun in Schwann cell cultures exposed to SAG for 48 hours. Densitometry of these blots is in (H). (Veh: DMSO vehicle). Data normalized to vehicle. One-way ANOVA with Dunnet’s test; *p<0.05, **p<0.005. n=6. (I) Representative Western blot showing c-Jun in Schwann cell cultures exposed to purmorphamine for 48 hours. Densitometry of these blots is in (J). Data normalized to vehicle. One-way ANOVA with Dunnet’s test; *p<0.05. n=3. (K) Representative immunofluorescence images showing increased c-Jun labeling of Sox10 positive Schwann cell nuclei after 48 hours incubation with SAG compared to vehicle (DMSO). Scale bar: 100μm. (L,M) qPCR showing mRNA expression of (L) BDNF *p=0.0314 and (M) GDNF *p=0.0382 in Schwann cell cultures incubated for 48 hours with SAG. Data normalized to vehicle. Unpaired Student’s t-tests. n=4 for each condition. (N) Differential interference contrast (DIC) microscopy showing changes in Schwann cell morphology after 48 hours incubation with purmorphamine (DMSO vehicle). Scale bar: 50μm. Graphs depict changes in cell area, roundness and length/√area following incubation with purmorphamine, demonstrating enhancement of elongated morphology. One-way ANOVAs with Tukey’s multiple comparison test ***p<0.001. n= 3, each experiment involving measurement of 100 cells per condition. (O) Representative Western blots showing c-Jun and phosphorylated c-Jun in cultured Schwann cells after 48 hours incubation with cyclopamine alone (DMSO vehicle). (P,Q) Densitometry of these blots. Data normalized to vehicle. One-way ANOVA with Dunnet’s multiple comparisons test; **p<0.005 in (P); **p<0.005, ***p<0.001 in (Q). n=3. All numerical data represented as means ± SEM.

After injury, Shh-dependent enhancement of c-Jun is likely to be mediated by Shh derived from Schwann cells, which are the major source of Shh in injured nerves. In support of such an autocrine hedgehog signaling loop, cyclopamine alone, which blocks cellular responses to Shh, down-regulated c-Jun protein and sharply suppressed c-Jun phosphorylation in cultured Schwann cells (Fig.9 O-Q).

Together these observations show that Shh promotes Schwann cell c-Jun expression in vitro and in vivo, and support the idea that injury triggers an autocrine Shh signaling loop to elevate c-Jun in repair cells.

## DISCUSSION

The present results indicate that reduced expression of c-Jun is an important factor in the repair cell failures seen during aging and chronic denervation. In both situations, Schwann cells of injured nerves fail to achieve or maintain high c-Jun levels, and in both cases, correction of c-Jun expression restores regeneration deficits. This highlights the importance of c-Jun in the function of repair Schwann cells, provides a common molecular link between two apparently unrelated problems in nerve repair, and points to manipulation of c-Jun regulated pathways as a potential route for improving the outcome of nerve injuries.

By using neuron back-filling this study provides a direct quantitative measure of neuronal regeneration capacity in vivo and how this is controlled by Schwann cell c-Jun levels. It also opens new questions that remain to be investigated. In particular, to what extent does c-Jun in Schwann cells determine other factors that are also important for repair. This includes the length of time neurons are able to sustain axon growth after injury, sprouting, axonal misrouting, targeting and synapse re-formation.

A previously identified gene set regulated by c-Jun in injured nerves (Arthur-Farraj et al. 2012), was found to be highly enriched among the genes affected by aging or chronic denervation in WT mice. The expression of a small group of genes was also positively correlated both with c-Jun levels and regeneration, suggesting that their role in Schwann cells or other cells in the nerve merits further study. In aging mice this encompasses Aqp5, Gpr37L1, Igfbp2 and Olig1, all of which have been studied in glial cells. Igfbp2 promotes phosphorylation of Akt, a pathway that is linked to Schwann cell proliferation and differentiation (reviewed in Ma et al. 2015; Boerboom et al. 2017; Jessen and Arthur-Farraj 2019). Gpr37L1 is a receptor for prosaposin and prosapeptide (Meyer et al. 2013). In Schwann cells prosapeptide phosphorylates MAPK (Hiraiwa et al. 1997) and prosaposin is secreted after nerve injury, facilitating regeneration (Hiraiwa et al. 1999). In experiments on chronic denervation this gene group encompasses Cxcl5, Egfl8, Gas2I3, Megf10 and Pcdh20. All of these genes were previously shown to be up-regulated in Schwann cells after injury (Zhang et al. 2011; Tanaka et al. 2013; Weiss et al. 2016; reviewed in Ma et al. 2016; Brosius Lutz et al. 2017). Cxcl5 activates STAT3 (Zhang et al. 2011), a transcription factor that we have shown to be important for maintaining repair cells during chronic denervation (Benito et al. 2017). Gas2I3 has a role in the cell cycle, and Megf10 in phagocytosis (Wolter et al. 2012; Chung et al. 2013).

Since c-Jun levels in injured nerves are a major determinant of effective repair, it is important to identify signals that control c-Jun expression. The present results suggest that Shh has a role in this process. In injured nerves of Shh cKO mice, there is reduced c-Jun activation and diminished Schwann cell expression of the c-Jun target p75NTR. In purified Schwann cells, application of Shh elevates c-Jun, while cyclopamine alone suppresses c-Jun. Shh also promotes Schwann cell elongation and is substantially reduced during chronic denervation. Previous work also implicates Shh signaling in repair. Shh is up-regulated in Schwann cells after injury (Hashimoto et al. 2008; Arthur-Farraj et al. 2012), and exposure to Shh improves nerve regeneration in various settings (Pepinsky et al. 2002; Bond et al. 2013; Martinez et al. 2015). Inhibition of Shh signaling reduces Schwann cell expression of BDNF, motor neuron survival after injury and axon regeneration (Hashimoto et al. 2008; Yamada et al. 2020), and a molecular link between Shh signaling and Jun activation has been established in various cell lines (Laner-Plamberger et al. 2009; Kudo et al. 2012). Further in vivo experiments using Shh cKO mice as well as Shh agonists and antagonists are needed to conclusively determine the involvement of Shh in regeneration. At present, however, the data presented here and previous work are consistent with the existence of an autocrine Shh signaling loop activated by injury to promote expression of c-Jun and the repair cell phenotype.

In c-Jun OE/+ mice, we considered whether restoration of c-Jun levels altered cell numbers, thus promoting regeneration. In aging mice, the results appear to exclude this, since cell numbers in the mutant and the WT are similar. During chronic denervation, Schwann cell numbers remain constant in c-Jun OE/+mice, but fall by about 30% in the WT. Since there is now evidence that Schwann cell proliferation may not be essential for regeneration, contrary to common assumptions, the relationship between cell numbers and repair is currently unclear (Kim et al. 2000; Atanasoski et al. 2001; Yang et al. 2008; for discussion see Jessen and Mirsky 2019). Even in WT mice, cell number after chronic denervation remains nearly twice that in uninjured nerves. It is therefore unlikely that the changes in Schwann cell numbers are the key reason for the reduced regeneration support provided by 10-week cut WT stumps, or the increase in support provided by 10 week cut c-Jun OE/+ stumps.

The degree of reduction in transverse nerve area after chronic denervation could also affect repair. However, the area of 10 week cut c-Jun OE/+ nerves, while increased compared to 10 week cut WT, remains ~50% smaller than that of 2-week cut WT nerves. Nevertheless, regeneration through these nerves is similar. The relationship between nerve area and regeneration in these experiments may therefore not be straightforward.

During longer denervation times, cell loss and nerve shrinking will increasingly impede repair. These slow, atrophic changes, likely involving regulation of cell death and proliferation, have not been extensively studied, although STAT3 has recently be implicated in the long term maintenance of repair cells (Benito et al. 2017). Previously, c-Jun was shown to influence both apoptosis and proliferation in repair Schwann cells (Parkinson et al. 2001; 2004; 2008), but the particular way in which c-Jun levels determine nerve atrophy remains to be determined.

It has become clear that the injury-induced reprogramming of Schwann cells to cells specialized to support nerve regeneration is regulated by dedicated mechanisms, including c-Jun, STAT3, merlin and H3K27 trimethylation-related epigenetic controls, that operate selectively in repair cells, and have a relatively minor or undetectable function in Schwann cell development (reviewed in Jessen and Mirsky 2019). The present work provides evidence that an impairment of one of these mechanisms, c-Jun, is a major contributor to two major categories of regeneration failure, aging and chronic denervation. It will be important to extend this study to other regulators of repair cells as a basis for developing molecular interventions for promoting repair in the PNS.

## METHODS

### Transgenic mice

Animal experiments conformed to UK Home Office guidelines under the supervision of University College London (UCL) Biological Services. Mice were generated to overexpress c-Jun selectively in Schwann cells as described (Fazal et al. 2017). Briefly, female *R26c-Junstopf* mice carrying a lox-P flanked STOP cassette in front of a CAG promoter-driven c-Jun cDNA in the ROSA26 locus, were crossed with male *P0Cre+/-* mice (Feltri et al. 1999). This generated *P0-Cre+/-/R26c-Junstopff/+* mice, referred to here as c-Jun OE/+ mice. *P0-Cre*-/- littermates were used as controls. Shh-floxed mice, referred to as Shh cKO mice, carrying loxP sites flanking exon 2 of the Shh gene were obtained from the Jackson Laboratory (Jax®, stock# 004293), and bred to P0cre mice (Feltri et al., 1999). Experiments used mice of either sex on the C57BL/6 background.

### Genotyping

DNA was extracted from ear notches or tail tips using the Hot Sodium Hydroxide and Tris method (HotSHot; (Truett et al. 2000). Tissue was incubated in HotSHot buffer (25 mM NaOH and 0.2 mM disodium EDTA, pH 12) at 95°C for 1h. The reaction was neutralized with neutralizing buffer (40 mM Tris-HCl, pH 5). DNA was then added to the PCR mastermix with primers for the P0-Cre transgene: 5’-GCTGGCCCAAATGTTGCTGG-3’ and 5’CCACCACCTCTCCATTGCAC-3’ (480 bp band).

### Surgery

For short-term time points (<1 week) and crushes, the sciatic nerve was exposed and cut or crushed (at 3 rotation angles) at the sciatic notch. For chronic denervation (>1 week), the sciatic nerve was cut and the proximal stump was reflected back and sutured into muscle to prevent regeneration. The nerve distal to the injury was excised for analysis at various time points. Contralateral uninjured sciatic nerves served as controls. To examine regeneration into distal stumps, the nerve branches of the sciatic nerve were individually separated. The tibial nerve was cut and both proximal and distal stumps were reflected and sutured into muscle. Either immediately or following 10 weeks of chronic denervation, the distal tibial nerve stump was cut from the muscle and sutured to the freshly transected common peroneal nerve.

### Retrograde labelling with Fluorogold

To examine regeneration, following nerve crush or repair, the nerve was cut distal to the original injury site and exposed to 4% Fluorogold for 1hour (Catapano et al. 2016). The spinal cord and L4 DRG were removed following perfusion 1 week post-labeling. Labeled cells in all of the spinal cord sections (50*μ*m) were counted and the Abercrombie correction was applied to compensate for double counting (Abercrombie 1946). To avoid double counting, cells in every fifth DRG section (20*μ*m) were counted.

### Schwann cell cultures

Rat Schwann cells were cultured as described (Brockes et al. 1979). Briefly, sciatic nerves and brachial plexuses were digested enzymatically with collagenase and trypsin and cultured on laminin and PLL coated plates in DMEM, 2% FBS, 10ng/ml NRG-1, 2*μ*M forskolin and penicillin/streptomycin. Under experimental conditions, cultures were maintained in defined medium (DMEM and Ham’s F12 (1:1), transferrin (100 pg/ml), progesterone (60 ng/ml), putrescine (16 pg/ml), insulin (5 *μ*g/ml), thyroxine (0.4 *μ*g/ml), selenium (160 ng/ml), triiodothyronine (10.1 ng/ml), dexamethasone (38 ng/ml), glucose (7.9 mg/ml), bovine serum albumin (0.3 mg/ml), penicillin (100 IU/ml), streptomycin (100 IU/ml), and glutamine (2 mM) with 0.5% serum (Jessen et al. 1994; Meier et al. 1999).

### Antibodies

Immunofluorescence antibodies: c-Jun (Cell Signaling Technology, rabbit 1:800), Sox10 (R&D Systems, goat 1:100), CGRP (Peninsula, rabbit 1:1000), Neurofilament (Abcam, rabbit 1:1000) donkey anti-goat IgG (H+L) Alexa Fluor 488 conjugate (Invitrogen, 1:1000), Cy3 donkey anti-rabbit IgG (H+L) (Jackson Immunoresearch, 1:500).

Antibodies used for Western blotting: c-Jun (Cell Signaling Technology, rabbit 1:1000), p75 NTR (Millipore, rabbit 1:1000), serine 63 phosphorylated c-Jun (Cell Signaling Technology, rabbit 1:1000), GAPDH (Sigma-Aldrich, rabbit 1:5000), calnexin (Enzo Life Sciences, rabbit 1:1000), anti-rabbit IgG, HRP-linked (Cell Signaling Technology, 1:2000).

### Immunofluorescence

For immunofluorescence experiments on cultured cells, 5000 Schwann cells were plated in a 35μl drop on a PLL laminin coated coverslip. Cells were topped up with defined medium after 24 hours. At the experimental end point, cells were washed 2x with 1x PBS. Cells were fixed with 4% PFA for 10 minutes. Cells were then washed for 5 minutes in 1x PBS. Fresh PBS was added to the wells and the lid was parafilm sealed. Dishes were stored at 4°C until use.

Nerve samples were fresh frozen during embedding in OCT. Cryosections were cut at 10μm. Sections were fixed in 100% acetone (Sox10/c-Jun double-labeling, 10 minutes at −20°C) or 4% PFA (10 minutes at room temperature).

For immunofluorescence, all samples were washed 3x in 1x PBS and blocked in 5% donkey serum, 1% BSA, 0.3% Triton X-100 in PBS. Samples were incubated with primary antibodies in blocking solution overnight at 4°C. Sox10/c-Jun double-labeling was performed overnight at room temperature. Samples were washed and incubated with secondary antibodies and DAPI to identify cell nuclei (Thermo Fisher Scientific, 1:40,000) in PBS for 1 hour at room temperature. Samples were mounted in fluorescent mounting medium (Citifluor).

Images were taken on a Nikon Labophot 2 fluorescence microscope. Cell counts were performed in ImageJ or directly from the microscope. Comparable images have been equally adjusted for brightness/contrast. In some cases (Figs 1, 3 and 4), images of whole nerve profiles have been generated by stitching together multiple images.

### Western Blotting

Nerves were dissected and snap frozen in liquid nitrogen. For protein extraction, nerves were placed in 2ml graduated skirted tubes with 9 10B lysing beads with 75ml lysis buffer (1M Tris-HCl pH 8, 5M NaCl, 20% Triton X-100, 5mM EDTA) and homogenized using a Fastprep fp120 homogeniser. Samples were run twice at speed 6 for 45 seconds. Lysates were then centrifuged at 13000rpm for 2 minutes at 4°C to pellet the debris. The supernatant was transferred to a new 1.5ml Eppendorf tube and centrifuged at 13000rpm for 2 minutes at 4°C. The supernatant was transferred to a new 1.5ml Eppendorf tube and the protein extract was stored at −80°C.

For protein studies on cultured cells, 1 x 10^6^ purified Schwann cells were plated in a 35mm dish in defined medium for 48 hours. At the time of extraction, the cultures were washed 2x with 1x PBS and incubated with 100μl cell lysis buffer (T-PER Tissue Protein Extraction Reagent, Halt protease & phosphatase inhibitor cocktail (1:100) Thermo Fisher Scientific). Cells were physically detached from dishes using a cell scraper. The cell lysate was collected and kept on ice in a 1.5ml Eppendorf tube. Lysate was spun for 2 minutes at 1000rpm to pellet the debris. The supernatant was transferred to a fresh Eppendorf tube and spun for a further 2 minutes at 1000rpm. The supernatant was transferred to a new 1.5ml Eppendorf tube and stored at −80°C until use.

Protein was diluted in the appropriate lysis buffer and 5x Laemmli buffer at a working concentration of 1x. Samples were heated to 95°C for 5 minutes to denature the protein. 10μg protein was loaded per well on 8% acrylamide gels with prestained standard molecular weight markers (PageRuler prestained protein ladder; Thermo Fisher Scientific) and run at 60mV for 3 hours using the mini Protean II gel electrophoresis apparatus (Bio-Rad Laboratories). Protein was transferred to a nitrocellulose membrane (Hybond ECL; Amersham Biosciences) using a semi-dry transfer system (Bio-Rad Laboratories) at 25mV for 45 minutes. Membranes were briefly stained with Ponceau S (Sigma Aldrich) to determine that the transfer has been successful and that equal levels of protein had been loaded in the gel. Membranes were briefly washed in ddH2O to remove excess Ponceau and blocked in 5% milk/TBS-T for 1 hour with shaking at room temperature. Membranes were then incubated with appropriate antibodies in heat sealable polyethylene bags and were incubated overnight at 4°C on a rotatory wheel. Membranes were washed 3x for 10 minutes in 1x TBS-T then incubated with the appropriate secondary antibody in heat sealable polyethylene bags, rotating for 1 hour at room temperature. Membranes were washed 3x for 10 minutes in 1x TBS-T before developing. For development of GAPDH, membranes were incubated with ECL (Amersham) for 1 minute and developed on a Bio-Rad Chemidoc machine. For the development of all other antibodies, membranes were incubated with ECL prime (Amersham) for 5 minutes then developed. Membranes were automatically exposed to prevent saturation. Blots were analysed and densitometric quantification was performed using Bio-Rad Imagelab. Protein levels were determined by normalising the protein of interest against the house keeping protein (GAPDH or calnexin). All blots were then normalized to one sample (e.g 1 week after injury, control cells) to account for any difference between each blot. Each experiment was performed at least 3 times with fresh samples. Representative images are shown.

### Electron microscopy

Nerves were fixed in 2.5% glutaraldehyde/2% paraformaldehyde in 0.1 M cacodylate buffer, pH 7.4, overnight at 4°C. Post-fixation in 1% OsO_4_ was performed before nerves were embedded in Agar 100 epoxy resin. Transverse ultrathin sections from adult (P60) or aged (P300) tibial nerves or from injured distal stumps of adult sciatic nerves at various times after injury were taken 5mm from the sciatic notch and mounted on film (no grid bars). Images were examined using a Jeol 1010 electron microscope with a Gatan camera and software. Images were examined and photographed at 8000× or 15,000x. The nerve area was measured from photographs taken at 200× magnification. Schwann cells and macrophages and fibroblasts were identified by standard ultrastructural criteria (e.g. Reichert et al. 1994). Schwann cell, macrophage and fibroblast nuclei were counted in every field, or every second, third or fourth field, depending on the size of the nerve, and multiplied by the number of fields to generate totals.

### qPCR

RNA from rat Schwann cell cultures or mouse nerve tissue was extracted using an RNeasy Micro Extraction Kit (Qiagen). RNA quality and concentration was determined after extraction using a nanodrop 2000 machine (Thermo). 1μg of RNA was converted to cDNA using SuperScriptTM II Reverse Transcriptase (Invitrogen) as per the manufacturer’s instructions. Samples were run with PrecisionPLUS qPCR Mastermix with SYBR Green (Primerdesign) with primers as described in Benito et al. (2017). Ct values were normalized to housekeeping gene expression (*GAPDH and calnexin*).

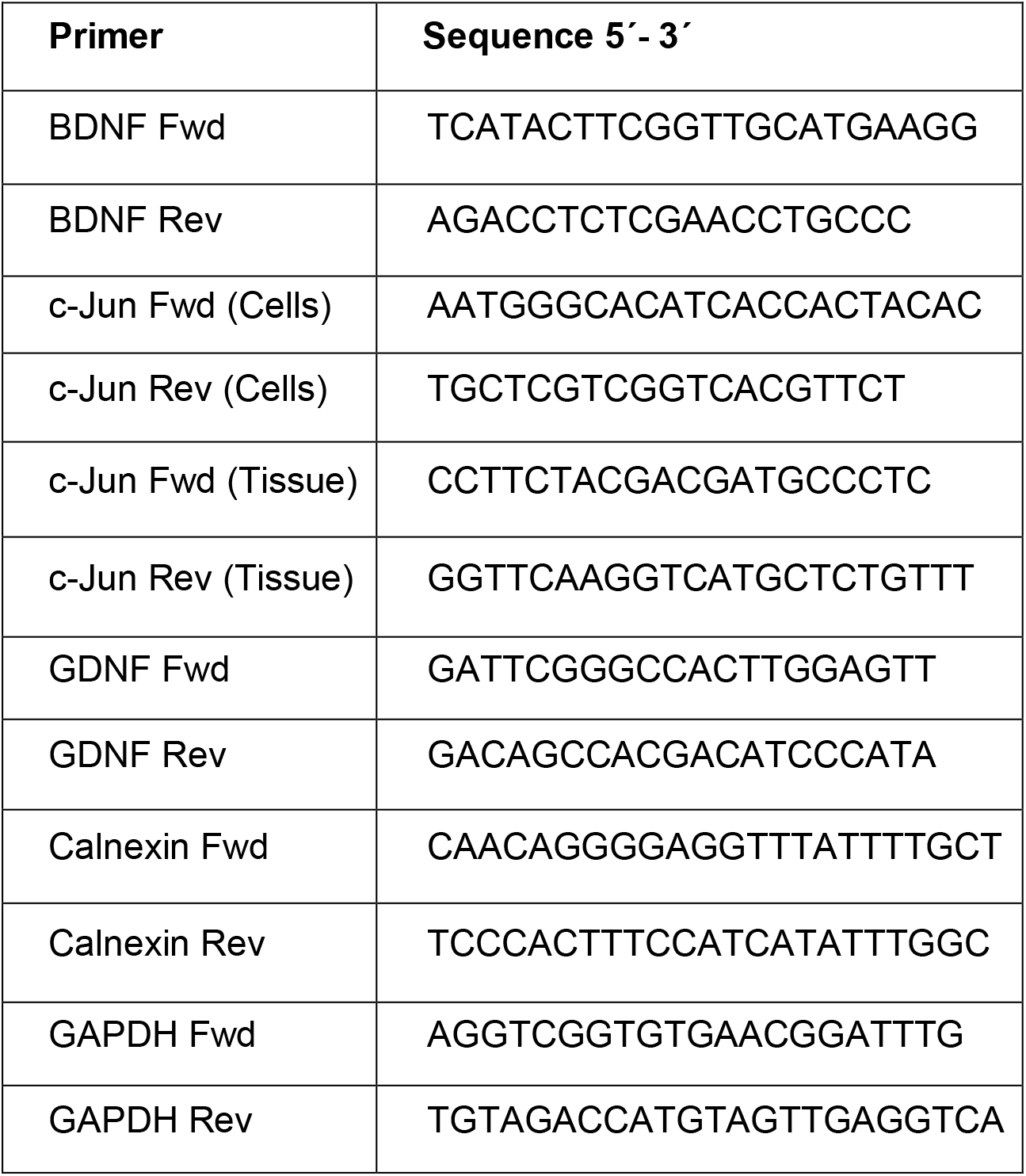

### Statistical analysis

Results are expressed as mean ± SEM. Statistical significance was estimated by Student’s *t* test, one-way ANOVA or two-way ANOVA with appropriate post hoc tests. A *p* value <0.05 was considered as statistically significant. Statistical analysis was performed using GraphPad software.

### Library preparation

RNA was extracted using a RNeasy lipid tissue kit with an in column DNase step (Qiagen). Chronically denervated and uninjured nerves were pooled, 2 per n. Acutely denervated nerves were not pooled using 1 nerve per n. The library was prepared using the Kapa mRNA Hyper Prep kit (Roche) with 100ng RNA and 15 cycles of PCR enrichment. The assay is (first) stranded (dUTP method).

### Sequencing

Sequencing was performed in a pooled NextSeq 500 run using a 43bp paired end protocol (plus a 6bp index read). Sequencing reads (in fastq format) were aligned to the hg38 reference sequence using STAR v2.5.3 (Dobin et al. 2013). Samtools version 1.2 and Picard tools version 1.140 were used to process alignments (Li et al. 2009)’ Aligned reads were filtered for mapq _ 4 i.e. uniquely mapping reads, and putative PCR duplicates were removed. featureCounts was used to perform read summarization (Liao et al. 2014).

### Data Analysis

Expression analysis was carried out using R version 3.5.1. Differential gene expression was analysed using edgeR (Robinson et al. 2010). Genes with both an absolute log2 fold change > 2.0 and FDR < 0.05 were identified as being significantly differentially expressed. Principal Component Analysis (PCA) showed that injury status was the dominant source of variation in both data sets (Fig. 7 Figure Supplement 1 A; Fig. 8 Figure Supplement 1A). Enrichment of c-Jun regulated genes was investigated using fisher one-sided exact tests and gene set enrichment analysis (GSEA; Subramanian et al. 2005). Gene ontology (GO) analysis was used to examine gene enrichment of all significantly differentiated genes using the PANTHER classification system (Mi et al. 2013).

## Acknowledgements

We are grateful to Jane Pendjiky for help with imaging.

## SUPPORTING FILES

**Supplementary Table 1: All significantly regulated genes in the tibial nerve of WT mice during aging and chronic denervation.** (A, B) Genes expressed at higher levels in aged mice under the conditions indicated are in blue (top), while genes with reduced expression in aged mice are in red (bottom). (C) Genes that respond more strongly to injury (3 day cut vs hUI) in aged mice are in blue (top), while genes with weaker injury response are in red (bottom). (D) The 303 genes that are significantly expressed in young WT vs UI and aged c-Jun OE/+ vs UI 3 days after injury. (E) Genes expressed at higher levels after chronic denervation are in blue (top), while genes with reduced expression are in red (bottom). (F) The 227 genes that are significantly expressed in WT 1 week vs UI and c-Jun OE/+ 10 week vs UI.

**Supplementary Table 2: 138 genes regulated by c-Jun in injured nerves derived from Arthur-Farraj et al. (2012).** Blue (top) indicates genes expressed at higher levels in cut nerves of WT mice compared with nerves of mice with conditional c-Jun inactivation selectively in Schwann cells (92 genes), while red (bottom) indicates genes expressed at lower levels in cut WT nerves compared to c-Jun mutant nerves (46 genes).

## REFERENCES

Abercrombie, M. 1946. Estimation of nuclear population from microtome sections. The Anatomical record 94, pp. 239–247.

Allodi, I., Udina, E. and Navarro, X. 2012. Specificity of peripheral nerve regeneration: interactions at the axon level. Progress in Neurobiology 98, pp. 16–37.

Arthur-Farraj, P.J., Claire C Morgan, C.C., Adamowicz, M., Gomez-Sanchez, J.A., Fazal, S.V., Beucher, A., Razzaghi, B., Mirsky, R., Jessen, K.R., Aitman, T.J. (2017). Changes in the Coding and Non-coding Transcriptome and DNA Methylome That Define the Schwann Cell Repair Phenotype After Nerve Injury. Cell Reports 20(11), pp. 2719–2734.

Arthur-Farraj, P.J., Latouche, M., Wilton, D.K., Quintes, S., Chabrol, E., Banerjee, A., Woodhoo, A., Jenkins, B., Rahman, M., Turmaine, M., Wicher, G.K., Mitter, R., Greensmith, L., Behrens, A., Raivich, G., Mirsky, R. and Jessen, K.R. 2012. c-Jun reprograms Schwann cells of injured nerves to generate a repair cell essential for regeneration. Neuron 75(4), pp. 633–647.

Atanasoski, S., Shumas, S., Dickson, C., Scherer, S.S. and Suter, U. 2001. Differential cyclin D1 requirements of proliferating Schwann cells during development and after injury. Molecular and Cellular Neurosciences 18(6), pp. 581–592.

Benito, C., Davis, C.M., Gomez-Sanchez, J.A., Turmaine, M., Meijer, D., Poli, V., Mirsky, R. and Jessen, K.R. 2017. STAT3 Controls the Long-Term Survival and Phenotype of Repair Schwann Cells during Nerve Regeneration. The Journal of Neuroscience 37(16), pp. 4255–4269.

Boerboom, A., Dion, V., Chariot, A. and Franzen, R. 2017. Molecular mechanisms involved in schwann cell plasticity. Frontiers in Molecular Neuroscience 10, p. 38.

Bond, C.W., Angeloni, N., Harrington, D., Stupp, S. and Podlasek, C.A. 2013. Sonic Hedgehog regulates brain-derived neurotrophic factor in normal and regenerating cavernous nerves. The Journal of Sexual Medicine 10(3), pp. 730–737.

Boyd, J. G., and Gordon, T. (2003). Neurotrophic factors and their receptors in axonal regeneration and functional recovery after peripheral nerve injury. Mol. Neurobiol. 27, pp. 277–324. doi: 10.1385/mn:27:3:277

Boyd, J.G. and Gordon, T. 2003. Glial cell line-derived neurotrophic factor and brain-derived neurotrophic factor sustain the axonal regeneration of chronically axotomized motoneurons in vivo. Experimental Neurology 183(2), pp. 610–619.

Brockes, J.P., Fields, K.L. and Raff, M.C. 1979. Studies on cultured rat Schwann cells. I. Establishment of purified populations from cultures of peripheral nerve. Brain Research 165(1), pp. 105–118.

Brosius Lutz, A., Chung, W.-S., Sloan, S.A., Carson, G.A., Zhou, L., Lovelett, E., Posada, S., Zuchero, J.B. and Barres, B.A. 2017. Schwann cells use TAM receptor-mediated phagocytosis in addition to autophagy to clear myelin in a mouse model of nerve injury. Proceedings of the National Academy of Sciences of the United States of America 114(38), pp. E8072–E8080.

Brushart, T. M., Aspalter, M., Griffin, J. W., Redett, R., Hameed, H., Zhou, C., Wright M, Vyas A, Höke A. 2013. Schwann cell phenotype is regulated by axon modality and central-peripheral location and persists in vitro. Experimental Neurology 247, pp. 272–281.

Büttner, R., Schulz, A., Reuter, M., Akula, A.K., Mindos, T., Carlstedt, A., Riecken, L.B., Baader, S.L., Bauer, R. and Morrison, H. 2018. Inflammaging impairs peripheral nerve maintenance and regeneration. Aging Cell 17(6), p. e12833.

Catapano, J., Willand, M.P., Zhang, J.J., Scholl, D., Gordon, T. and Borschel, G.H. 2016. Retrograde labeling of regenerating motor and sensory neurons using silicone caps. Journal of Neuroscience Methods 259, pp. 122–128.

Chen, Z.L., Yu, W.M. and Strickland, S. 2007. Peripheral regeneration. Annual Review of Neuroscience. 30, pp. 209–33.

Chung, W.-S., Clarke, L.E., Wang, G.X., Stafford, B.K., Sher, A., Chakraborty, C., Joung, J., Foo, L.C., Thompson, A., Chen, C., Smith, S.J. and Barres, B.A. 2013. Astrocytes mediate synapse elimination through MEGF10 and MERTK pathways. Nature 504(7480), pp. 394–400.

Clements, M.P., Byrne, E., Camarillo Guerro, L.F., Cattin, A., Zakka, L., Ashraf, A., Burden, J.J., Khadayate, S., Lloyd, A.C., Marguerat, S., Parrinello, S. 2017. The Wound Microenvironment Reprograms Schwann Cells to Invasive Mesenchymal-like Cells to Drive Peripheral Nerve Regeneration. Neuron 96(1), pp 98–114.e7

Cohen, D.E. and Melton, D. 2011. Turning straw into gold: directing cell fate for regenerative medicine. Nature Review in Genetics 12, pp. 243–252.

Dobin, A. et al., 2013. STAR: ultrafast universal RNA-seq aligner. Bioinformatics, 29(1), pp. 15–21.

Doron-Mandel, E., Fainzilber, M. and Terenzio, M. 2015. Growth control mechanisms in neuronal regeneration. FEBS Letters 589, pp. 1669–1677.

Eggers, R., Tannemaat, M.R., Ehlert, E.M. and Verhaagen, J. 2010. A spatio-temporal analysis of motoneuron survival, axonal regeneration and neurotrophic factor expression after lumbar ventral root avulsion and implantation. Experimental Neurology 223(1), pp. 207–220.

Eguizabal, C., Montserrat, N., Veiga A.. and Izpisua Belmonte, J.C. 2013. Dedifferentiation, transdifferentiation, and reprogramming: future directions in regenerative medicine. Seminars in. Reproductive Medicine 31, pp. 82–94.

Fawcett, J.W. and Verhaagen, J. 2018. Intrinsic Determinants of Axon Regeneration. Developmental Neurobiology 78(10), pp. 890–897.

Fazal, S.V., Gomez-Sanchez, J.A., Wagstaff, L.J., Musner, N., Otto, G., Janz, M., Mirsky, R. and Jessen, K.R. 2017. Graded Elevation of c-Jun in Schwann Cells In Vivo: Gene Dosage Determines Effects on Development, Remyelination, Tumorigenesis, and Hypomyelination. The Journal of Neuroscience 37(50), pp. 12297–12313.

Feltri, M.L., D’Antonio, M., Previtali, S., Fasolini, M., Messing, A. and Wrabetz, L. 1999. P0-Cre transgenic mice for inactivation of adhesion molecules in Schwann cells. Annals of the New York Academy of Sciences 883, pp. 116–123.

Fontana, X., Hristova, M., Da Costa, C., Patodia, S., Thei, L., Makwana, M., Spencer-Dene, B., Latouche, M., Mirsky, R., Jessen, K.R., Klein, R., Raivich, G., Behrens, A. 2012. c-Jun in Schwann cells promotes axonal regeneration and motoneuron survival via paracrine signalling. Journal of Cell Biology 198(1), pp. 127–141

Fu, S.Y. and Gordon, T. 1995. Contributing factors to poor functional recovery after delayed nerve repair: prolonged denervation. The Journal of Neuroscience 15(5 Pt 2), pp. 3886–3895.

Furey, M.J., Midha, R., Xu, Q.-G., Belkas, J. and Gordon, T. 2007. Prolonged target deprivation reduces the capacity of injured motoneurons to regenerate. Neurosurgery 60(4), pp. 723–732.

Gambarotta, G., Fregnan, F., Gnavi, S. and Perroteau, I. 2013. Neuregulin 1 role in Schwann cell regulation and potential applications to promote peripheral nerve regeneration. International Review of Neurobiology 108, pp. 223–256.

Glenn, T. D. and Talbot, W. S. 2013. Signals regulating myelination in peripheral nerves and the Schwann cell response to injury. Current Opininion in Neurobiology 23, pp. 1041–1048.

Gomez-Sanchez, J.A., Pilch, K.S., van der Lans, M., Fazal, S.V., Benito, C., Wagstaff, L.J., Mirsky, R. and Jessen, K.R. 2017. After nerve injury, lineage tracing shows that myelin and Remak Schwann cells elongate extensively and branch to form repair schwann cells, which shorten radically on remyelination. The Journal of Neuroscience 37(37), pp. 9086–9099.

Graciarena, M., Dambly-Chaudière, C. and Ghysen, A. 2014. Dynamics of axonal regeneration in adult and aging zebrafish reveal the promoting effect of a first lesion. Proceedings of the National Academy of Sciences of the United States of America 111(4), pp. 1610–1615.

Grothe, C., Haastert, K. and Jungnickel, J. 2006. Physiological function and putative therapeutic impact of the FGF-2 system in peripheral nerve regeneration—lessons from in vivo studies in mice and rats. Brain Research Reviews 51, pp. 293–299.

Hashimoto, M., Ishii, K., Nakamura, Y., Watabe, K., Kohsaka, S. and Akazawa, C. 2008. Neuroprotective effect of sonic hedgehog up-regulated in Schwann cells following sciatic nerve injury. Journal of Neurochemistry 107(4), pp. 918–927.

Hiraiwa, M., Campana, W.M., Mizisin, A.P., Mohiuddin, L. and O’Brien, J.S. 1999. Prosaposin: a myelinotrophic protein that promotes expression of myelin constituents and is secreted after nerve injury. Glia 26(4), pp. 353–360.

Hiraiwa, M., Taylor, E.M., Campana, W.M., Darin, S.J. and O’Brien, J.S. 1997. Cell death prevention, mitogen-activated protein kinase stimulation, and increased sulfatide concentrations in Schwann cells and oligodendrocytes by prosaposin and prosaptides. Proceedings of the National Academy of Sciences of the United States of America 94(9), pp. 4778–4781.

Höke, A. 2006. Mechanisms of disease: what factors limit the success of peripheral nerve regeneration in humans? Nature Clinical Practice Neurology 2, 448–454.

Höke, A., Gordon, T., Zochodne, D.W. and Sulaiman, O.A.R. 2002. A decline in glial cell-line-derived neurotrophic factor expression is associated with impaired regeneration after longterm Schwann cell denervation. Experimental Neurology 173(1), pp. 77–85.

Hsin-Pin, L., Oksuz, I., Hurley, E.., Wrabetz, L., Awatramani, R., 2015. Microprocessor Complex Subunit DiGeorge Syndrome Critical Region Gene 8 (Dgcr8) Is Required for Schwann Cell Myelination and Myelin Maintenance. Journal of Biological Chemistry, 290, pp. 24294–24307.

Huang, L., Quan, X., Liu, Z., Ma, T., Wu, Y., Ge, J., Zhu, S., Yang, Y., Liu, L., Sun, Z., Huang, J. and Luo, Z. 2015. c-Jun gene-modified Schwann cells: upregulating multiple neurotrophic factors and promoting neurite outgrowth. Tissue Engineering. Part A 21(7-8), pp. 1409–1421.

Huang, L., Xia, B., Shi, X., Gao, J., Yang, Y., Xu, F., Qi, F., Liang, C., Huang, J. and Luo, Z. 2019. Time-restricted release of multiple neurotrophic factors promotes axonal regeneration and functional recovery after peripheral nerve injury. The FASEB Journal, 33(7), pp.8600–8613.

Jessen, K.R. and Arthur-Farraj, P. 2019. Repair Schwann cell update: Adaptive reprogramming, EMT, and stemness in regenerating nerves. Glia 67(3), pp. 421–437.

Jessen, K.R., Brennan, A., Morgan, L., Mirsky, R., Kent, A., Hashimoto, Y. and Gavrilovic, J. 1994. The Schwann cell precursor and its fate: a study of cell death and differentiation during gliogenesis in rat embryonic nerves. Neuron 12(3), pp. 509–527.

Jessen, K.R. and Mirsky, R. 2016. The repair Schwann cell and its function in regenerating nerves. The Journal of Physiology 594(13), pp. 3521–3531.

Jessen, K.R. and Mirsky, R. 2019. The success and failure of the schwann cell response to nerve injury. Frontiers in Cellular Neuroscience 13, p. 33.

Jessen, K.R., Mirsky, R., Arthur-Farraj, P. 2015. The Role of Cell Plasticity in Tissue Repair: Adaptive Cellular Reprogramming. Developmental Cell 34, pp. 613–620

Jonsson, S., Wiberg, R., McGrath, A.M., Novikov, L.N., Wiberg, M., Novikova, L.N. and Kingham, P.J. 2013. Effect of delayed peripheral nerve repair on nerve regeneration, Schwann cell function and target muscle recovery. Plos One 8(2), p. e56484.

Kim, H.A., Pomeroy, S.L., Whoriskey, W., Pawlitzky, I., Benowitz, L.I., Sicinski, P., Stiles, C.D. and Roberts, T.M. 2000. A developmentally regulated switch directs regenerative growth of Schwann cells through cyclin D1. Neuron 26(2), pp. 405–416.

Kudo, K., Gavin, E., Das, S., Amable, L., Shevde, L.A. and Reed, E. 2012. Inhibition of Gli1 results in altered c-Jun activation, inhibition of cisplatin-induced upregulation of ERCC1, XPD and XRCC1, and inhibition of platinum–DNA adduct repair. Oncogene, 31(44), pp. 4718–4724.

Laner-Plamberger, S., Kaser, A., Paulischta, M., Hauser-Kronberger, C., Eichberger, T. and Frischauf, A. M. 2009. Cooperation between GLI and JUN enhances transcription of JUN and selected GLI target genes. Oncogene 28(13), pp. 1639–1651.

Li, H., Handsaker, B., Wysoker, A., Fennell, T., Ruan, J., Homer, N., Marth, G., Abecasis, G., Durbin, R. 2009. The Sequence Alignment/Map format and SAMtools. Bioinformatics, 25(16), pp. 2078–2079.

Liao, Y., Smyth, G.K. & Shi, W., 2014. featureCounts: an efficient general purpose program for assigning sequence reads to genomic features. Bioinformatics, 30(7), pp. 923–930.

Lu, Q.R., Yuk, D., Alberta, J.A., Zhu, Z., Pawlitzky, I., Chan, J., McMahon, A.P., Stiles, C.D., Rowitch, D.H. 2000. Sonic hedgehog--regulated oligodendrocyte lineage genes encoding bHLH proteins in the mammalian central nervous system. Neuron. 25, pp. 317–329.

Ma, K.H., Hung, H.A., Srinivasan, R., Xie, H., Orkin, S.H. and Svaren, J. 2015. Regulation of Peripheral Nerve Myelin Maintenance by Gene Repression through Polycomb Repressive Complex 2. The Journal of Neuroscience 35(22), pp. 8640–8652.

Ma, K.H., Hung, H.A. and Svaren, J. 2016. Epigenomic regulation of schwann cell reprogramming in peripheral nerve injury. The Journal of Neuroscience 36(35), pp. 9135–9147.

Martinez, J.A., Kobayashi, M., Krishnan, A., Webber, C., Christie, K., Guo, G., Singh, V. and Zochodne, D.W. 2015. Intrinsic facilitation of adult peripheral nerve regeneration by the Sonic hedgehog morphogen. Experimental Neurology 271, pp. 493–505.

Meier, C., Parmantier, E., Brennan, A., Mirsky, R. and Jessen, K.R. 1999. Developing Schwann cells acquire the ability to survive without, axons by establishing an autocrine circuit involving Insulin-like growth factor, neurotrophin-3, and Platelet-derived growth factor-BB. Journal of Neuroscience 19(10), pp. 3847–3859.

Melcangi, R.C., Magnaghi, V., Cavarretta, I., Riva, M.A., Piva, F. and Martini, L. 1998. Effects of steroid hormones on gene expression of glial markers in the central and peripheral nervous system: variations induced by aging. Experimental Gerontology 33(7-8), pp 827–36.

Melcangi, R.C, Magnaghi, V. and Martini L. 2000. Aging in peripheral nerves: regulation of myelin protein genes by steroid hormones. Progress in Neurobiology 60(3), pp. 291–308.

Meyer, R.C., Giddens, M.M., Schaefer, S.A. and Hall, R.A. 2013. GPR37 and GPR37L1 are receptors for the neuroprotective and glioprotective factors prosaptide and prosaposin. Proceedings of the National Academy of Sciences of the United States of America 110(23), pp. 9529–9534.

Mi, H., Muruganujan, A., Casagrande, J.T., and Thomas, .PD. 2013. Large-scale gene function analysis with the PANTHER classification system. Nature Protocols, 8(8), pp. 1551–1566.

Michalski, B., Bain, J.R. and Fahnestock, M. 2008. Long-term changes in neurotrophic factor expression in distal nerve stump following denervation and reinn ervation with motor or sensory nerve. Journal of Neurochemistry 105(4), pp. 1244–1252.

Nocera, G. and Jacob, C. 2020. Mechanisms of Schwann cell plasticity involved in peripheral nerve repair after injury. Cellular and Molecular Life Sciences.

Novikova, L., Novikov, L. and Kellerth, J.O. 1997. Persistent neuronal labeling by retrograde fluorescent tracers: a comparison between Fast Blue, Fluoro-Gold and various dextran conjugates. Journal of Neuroscience Methods 74(1), pp. 9–15.

Painter, M.W. 2017. Aging Schwann cells: mechanisms, implications, future directions. Current Opinion in Neurobiology 47, pp. 203–208.

Painter, M.W., Brosius Lutz, A., Cheng, Y.-C., Latremoliere, A., Duong, K., Miller, C.M., Posada, S., Cobos, E.J., Zhang, A.X., Wagers, A.J., Havton, L.A., Barres, B., Omura, T. and Woolf, C.J. 2014. Diminished Schwann cell repair responses underlie age-associated impaired axonal regeneration. Neuron 83(2), pp. 331–343.

Parkinson, D.B., Bhaskaran, A., Arthur-Farraj, P., Noon, L.A., Woodhoo, A., Lloyd, A.C., Feltri, M.L., Wrabetz, L., Behrens, A., Mirsky, R. and Jessen, K.R. 2008. c-Jun is a negative regulator of myelination. The Journal of Cell Biology 181(4), pp. 625–637.

Parkinson, D.B., Bhaskaran, A., Droggiti, A., Dickinson, S., D’Antonio, M., Mirsky, R. and Jessen, K.R. 2004. Krox-20 inhibits Jun-NH2-terminal kinase/c-Jun to control Schwann cell proliferation and death. The Journal of Cell Biology 164(3), pp. 385–394.

Parkinson, D.B., Dong, Z., Bunting, H., Whitfield, J., Meier, C., Marie, H., Mirsky, R. and Jessen, K.R., 2001. Transforming growth factor β (TGFβ) mediates Schwann cell death in vitro and in vivo: examination of c-Jun activation, interactions with survival signals, and the relationship of TGFβ-mediated death to Schwann cell differentiation. Journal of Neuroscience, 21(21), pp. 8572–8585.

Pepinsky, R.B., Shapiro, R.I., Wang, S., Chakraborty, A., Gill, A., Lepage, D.J., Wen, D., Rayhorn, P., Horan, G.S., Taylor, F.R. and Garber, E.A., 2002. Long-acting forms of sonic hedgehog with improved pharmacokinetic and pharmacodynamic properties are efficacious in a nerve injury model. Journal of pharmaceutical sciences, 91(2), pp. 371–387.

Pestronk, A., Drachman, D.B. and Griffin, J.W. 1980. Effects of aging on nerve sprouting and regeneration. Experimental Neurology 70(1), pp. 65–82.

Reichert, F., Saada, A. and Rotshenker, S. 1994. Peripheral nerve injury induces Schwann cells to express two macrophage phenotypes: phagocytosis and galagtose specific lectin MAC-2. Journal of Nueroscience 14, pp. 3231–3245.

Robinson, M.D., McCarthy, D.J. & Smyth, G.K., 2010. edgeR: a Bioconductor package for differential expression analysis of digital gene expression data. Bioinformatics, 26(1), pp. 139–140.

Ruijs, A.C.J., Jaquet, J., Kalmijn, S., Giele, H., Hovius, S.E.R. 2005. Median and ulnar nerve injuries: A meta-analysis of predictors of motor and sensory recovery after modern microsurgical nerve repair. Plastic Reconstructive Surgery 116(2), pp. 484–494.

Scheib, J. and Höke A. 2013. Advances in peripheral nerve regeneration. Nature Reviews Neurology 9(12), pp. 668–676.

Scheib, J.L. and Höke, A. 2016. An attenuated immune response by Schwann cells and macrophages inhibits nerve regeneration in aged rats. Neurobiology of Aging 45, pp. 1–9.

Subramanian, A., Tamayo, P., Mootha, V.K., Mukherjee, S., Ebert, B.L., Gillette, M.A., Paulovich, A., Pomeroy, S.L., Golub, T.R., Lander, E.S., and Mesirov, J.P. 2005. Gene set enrichment analysis: a knowledge-based approach for interpreting genome-wide expression profiles. Proceedings of the National Academy of Sciences of the United States of America, 102(43), pp. 15545–15550.

Sulaiman, O.A. and Gordon, T. 2000. Effects of short- and long-term Schwann cell denervation on peripheral nerve regeneration, myelination, and size. Glia 32(3), pp. 234–246.

Sulaiman, O. A. and Gordon, T. 2009. Role of chronic Schwann cell denervation in poor functional recovery after nerve injuries and experimental strategies to combat it. Neurosurgery 65, A105–A114.

Tanaka, K., Eskin, A., Chareyre, F., Jessen, W.J., Manent, J., Niwa-Kawakita, M., Chen, R., White, C.H., Vitte, J., Jaffer, Z.M., Nelson, S.F., Rubenstein, A.E. and Giovannini, M. 2013. Therapeutic potential of HSP90 inhibition for neurofibromatosis type 2. Clinical Cancer Research 19(14), pp. 3856–3870.

Tanaka, K. and Webster, H.D. 1991. Myelinated fiber regeneration after crush injury is retarded in sciatic nerves of aging mice. The Journal of Comparative Neurology 308(2), pp. 180–187.

Tanaka, K., Zhang, Q.L. and Webster, H.D. 1992. Myelinated fiber regeneration after sciatic nerve crush: morphometric observations in young adult and aging mice and the effects of macrophage suppression and conditioning lesions. Experimental Neurology 118(1), pp. 53–61.

Truett, G.E., Heeger, P., Mynatt, R.L., Truett, A.A., Walker, J.A. and Warman, M.L. 2000. Preparation of PCR-quality mouse genomic DNA with hot sodium hydroxide and tris (HotSHOT). Biotechniques 29(1), pp. 52–54.

Vaughan, D.W. 1992. Effects of advancing age on peripheral nerve regeneration. The Journal of Comparative Neurology 323(2), pp. 219–237.

Verdú, E., Ceballos, D., Vilches, J.J. and Navarro, X. 2000. Influence of aging on peripheral nerve function and regeneration. Journal of the Peripheral Nervous System 5(4), pp. 191–208.

Weiss, T., Taschner-Mandl, S., Bileck, A., Slany, A., Kromp, F., Rifatbegovic, F., Frech, C., Windhager, R., Kitzinger, H., Tzou, C.-H., Ambros, P.F., Gerner, C. and Ambros, I.M. 2016. Proteomics and transcriptomics of peripheral nerve tissue and cells unravel new aspects of the human Schwann cell repair phenotype. Glia 64(12), pp. 2133–2153.

Wilcox, M.B., Laranjeira, S.G., Eriksson, T.M., Jessen, K.R., Mirsky, R., Quick, T.J. and Phillips, J.B. 2020. Characterising cellular and molecular features of human peripheral nerve degeneration. Acta Neuropathologica Communications 8(1), p. 51.

Wolter, P., Schmitt, K., Fackler, M., Kremling, H., Probst, L., Hauser, S., Gruss, O.J. and Gaubatz, S. 2012. GAS2L3, a target gene of the DREAM complex, is required for proper cytokinesis and genomic stability. Journal of Cell Science 125(Pt 10), pp. 2393–2406.

Yamada, Y., Trakanant, S., Nihara, J., Kudo, T., Seo, K., Saeki, M., Kurose, M., Matsumaru, D., Maeda, T. and Ohazama, A. 2020. Gli3 is a Key Factor in the Schwann Cells from Both Intact and Injured Peripheral Nerves. Neuroscience 432, pp. 229–239.

Yang, D.P., Zhang, D.P., Mak, K.S., Bonder, D.E., Pomeroy, S.L. and Kim, H.A. 2008. Schwann cell proliferation during Wallerian degeneration is not necessary for regeneration and remyelination of the peripheral nerves: axon-dependent removal of newly generated Schwann cells by apoptosis. Molecular and Cellular Neurosciences 38(1), pp. 80–88.

You, S., Petrov, T., Chung, P.H. and Gordon, T. 1997. The expression of the low affinity nerve growth factor receptor in long-term denervated Schwann cells. Glia 20(2), pp. 87–100.

Zhang, H., Yang, R., Wang, Z., Lin, G., Lue, T.F. and Lin, C.-S. 2011. Adipose tissue-derived stem cells secrete CXCL5 cytokine with neurotrophic effects on cavernous nerve regeneration. The Journal of Sexual Medicine 8(2), pp. 437–446.

Zhou, Q., Wang, S., Anderson, D.J. 2000. Identification of a novel family of oligodendrocyte lineage-specific basic helix-loop-helix transcription factors. Neuron 25(2), pp. 331–343.

